# Two distinct phases of chloroplast biogenesis during de-etiolation in *Arabidopsis thaliana*

**DOI:** 10.1101/2020.08.30.274043

**Authors:** Rosa Pipitone, Simona Eicke, Barbara Pfister, Gaetan Glauser, Denis Falconet, Clarisse Uwizeye, Thibaut Pralon, Samuel Zeeman, Felix Kessler, Emilie Demarsy

## Abstract

Light triggers chloroplast differentiation whereby the etioplast transforms into a photosynthesizing chloroplast and the thylakoid rapidly emerges. However, the sequence of events during chloroplast differentiation remains poorly understood. Using Serial Block Face Scanning Electron Microscopy (SBF-SEM), we generated a series of chloroplast 3D reconstructions during differentiation, revealing chloroplast number and volume and the extent of envelope and thylakoid membrane surfaces. Furthermore, we used quantitative lipid and whole proteome data to complement the (ultra)structural data, providing a time-resolved, multi-dimensional description of chloroplast differentiation. This showed two distinct phases of chloroplast biogenesis: an initial photosynthesis-enabling ‘Structure Establishment Phase’ followed by a ‘Chloroplast Proliferation Phase’ during cell expansion. Moreover, these data detail thylakoid membrane expansion during de-etiolation at the seedling level and the relative contribution and differential regulation of proteins and lipids at each developmental stage. Altogether, we establish a roadmap for chloroplast differentiation, a critical process for plant photoautotrophic growth and survival.

## Introduction

Seedling development relies on successful chloroplast biogenesis, ensuring the transition from heterotrophic to autotrophic growth. Light is a crucial factor for chloroplast differentiation. For seeds that germinate in the light, chloroplasts may differentiate directly from proplastids present in cotyledons. However, as seeds most often germinate underneath soil, seedling development typically begins in darkness and follows a skotomorphogenic program called etiolation, characterized by rapid hypocotyl elongation and etioplast development. Light promotes seedling de-etiolation, which involves a series of morphological changes, such as cotyledon expansion, hypocotyl growth inhibition, and greening, that accompanies the onset of photosynthesis in chloroplasts. During de-etiolation, etioplast–chloroplast transition is thereby rapidly triggered by light following seedling emergence at the soil surface (Solymosi and Schoefs, 2010; Weier and Brown, 1970). A hallmark of chloroplast differentiation is the biogenesis of thylakoids, a network of internal membranes where the components of the photosynthetic electron transport chain assemble. Thylakoid biogenesis and the onset of photosynthesis rely on the concerted synthesis and coordinated assembly of lipids and proteins in both space and time.

The thylakoids harbor the photosynthetic electron transport chain, which is composed of three complexes: photosystem II (PSII), the cytochrome *b*_6_*f* complex (Cyt *b*_6_*f*), and photosystem I (PSI). Electron transfer between these complexes is facilitated by mobile electron carriers, specifically the low-molecular-weight, membrane-soluble plastoquinone (electron transfer from PSII to Cyt *b*_6_*f*) and the lumenal protein plastocyanin (electron transfer from Cyt *b*_6_*f* to PSI). Electron transfer leads to successive reduction and oxidation of electron transport chain components. The final reduction step catalyzed by ferredoxin-NADP(+) reductase (FNR) leads to NADPH production. Oxidation of water by PSII and of plastoquinone by Cyt *b*_6_*f* releases protons into the lumen, generating a proton gradient across the thylakoid membrane that drives the activity of the thylakoid-localized chloroplast ATP synthase complex. Each of the photosynthetic complexes consists of multiple subunits encoded by the plastid or nuclear genome. PSII and PSI have core complexes comprising 25–30 and 15 proteins, respectively (Amunts and Nelson, 2009; Caffarri et al., 2014). The antenna proteins from the Light Harvesting Complexes (LHC) surround the PSI and PSII core complexes contributing to the formation of supercomplexes. Cyt *b*_6_*f* is an eight-subunit dimeric complex. Each complex of the electron transport chain has a specific dimension, orientation, and location within the thylakoid membrane, occupying a defined surface, and their dimensions have been reported in several studies giving congruent results (Caffarri et al., 2014; Kurisu et al., 2003; Van Bezouwen et al., 2017). During de-etiolation, massive protein synthesis is required for assembly of the highly abundant photosynthetic complexes embedded in thylakoids. Chloroplast proteins encoded by the nuclear genome must be imported from the cytoplasm. The general chloroplast protein import machinery is composed of the multimeric complexes Translocon of Outer membrane Complex (TOC) and Translocon of Inner membrane Complex (TIC), and selective import is based on specific recognition of transit peptide sequences and TOC receptors (Agne and Kessler, 2010; Richardson and Schnell, 2019).

Reminiscent of their cyanobacterial origin, chloroplast membranes are composed mostly of glycolipids (mono- and di-galactosyldiacylglycerol; MGDG and DGDG) and are poor in phospholipids compared to other membranes in the cell (Bastien et al., 2016; Block et al., 1983; Kobayashi, 2016). Galactolipids comprise a glycerol backbone esterified to contain a single (MGDG) or double (DGDG) galactose units at the *sn*1 position and two fatty acid chains at the *sn*2 and *sn*3 positions. In addition to the number of galactose units at *sn*1, galactolipids also differ by the length and degrees of saturation of the fatty acid chains. In some species, including Arabidopsis, galactolipid synthesis relies on two different pathways, defined as the eukaryotic and prokaryotic pathway depending on the organellar origin of the diacylglycerol precursor. The eukaryotic pathway requires the import of diacyl-glycerol (DAG) synthesized in the endoplasmic reticulum (ER) into the plastids and is referred to as the ER pathway, whereas the prokaryotic pathway is entirely restricted to the plastid (PL) and is referred to as the PL pathway (Ohlrogge and Browse, 1995). As signatures, ER pathway–derived galactolipids harbor an 18-carbon chain whereas PL pathway–derived galactolipids harbor a 16-carbon chain at the *sn*2 position. In addition to constituting the lipid bilayer, galactolipids are integral components of photosystems and thereby contribute to photochemistry and photoprotection (Aronsson et al., 2008; Kobayashi, 2016). Thylakoids also contain neutral lipids such as chlorophyll, carotenoids, tocopherols, and plastoquinone. These may exist freely or be associated with the photosynthetic complexes, having either a direct role in photosynthesis (chlorophyll, carotenoids, plastoquinone) or participating indirectly in the optimization of light usage and/or mitigation of potentially damaging effects (tocopherols in addition to carotenoids and plastoquinone) (Hashimoto et al., 2003; Van Wijk and Kessler, 2017).

Past studies used conventional electron microscopy to first describe the architecture of the thylakoid membrane network. Based on these 2D observations, researchers proposed that plant thylakoid membranes are organised as single lamellae connected to appressed multi-lamellar regions called grana. How these lamellae are interconnected was revealed only later following the development of 3D electron microscopic techniques. Tremendous technological progress in the field of electron microscopy has been made recently, leading to improved descriptions of chloroplast ultrastructure (Daum et al., 2010; Daum and Kühlbrandt, 2011). Electron tomography substantially improved our comprehension of the 3D organisation of the thylakoid network in chloroplasts at different developmental stages and in different photosynthetic organisms, including Arabidopsis (Austin and Staehelin, 2011; Liang et al., 2018), Chlamydomonas (Engel et al., 2015), runner bean (Kowalewska et al., 2016), and *Phaeodactylum tricornutum* (Flori et al., 2017). Electron tomography also provided quantitative information on thylakoid structure such as the thylakoid layer number within the grana stack and the thickness of the stacking repeat distance of grana membrane (Daum et al., 2010; Kirchhoff et al., 2011). These quantitative data allowed a greater understanding of the spatial organisation of the thylakoid membrane in relation to the embedded photosynthetic complexes (Wietrzynski et al., 2020). Although electron tomography offers extraordinary resolution at the nanometer level, its main drawback is a limit to the volume of the observation, enabling only a partial 3D reconstruction of a chloroplast. SBF-SEM technology allows a much larger volume to be studied and reconstructed in 3D to show cellular organisation (Peddie and Collinson, 2014; Pinali and Kitmitto, 2014).

In combination with electron microscopy, biochemical fractionation of thylakoids has revealed differential lipid and protein compositions of the grana and the stroma lamellae. The grana are enriched in DGDG and PSII whereas the stroma lamellae are enriched in MGDG, Cyt *b6/f*, and PSI (Demé et al., 2014; Koochak et al., 2019; Tomizioli et al., 2014; Wietrzynski et al., 2020). Changes in lipid and protein compositions during etioplast–chloroplast transition are tightly linked to the thylakoid architecture. In particular, changes in MGDG to DGDG ratio are correlated with the transition from prolamellar body (PLB) and prothylakoid (PT) structures (tubular membrane) to thylakoid membranes (lamellar structure) (Bottier et al., 2007; Demé et al., 2014; Mazur et al., 2019).

Individual studies have provided much insight regarding specific dynamics of the soluble chloroplast proteome, the chloroplast transcriptome, photosynthesis-related protein accumulation and photosynthetic activity, chloroplast lipids, and changes in thylakoid architecture (Armarego-Marriott et al., 2019; Dubreuil et al., 2018; Kleffmann et al., 2007; Kowalewska et al., 2016; Liang et al., 2018; Rudowska et al., 2012). However, these studies were mostly qualitative, focused on one or two aspects, and were performed in different model organisms. Therefore, chemical data related to thylakoid biogenesis remain sparse and quantitative information is rare. Here, we present a systems-level study that integrates quantitative information on ultrastructural changes of the thylakoids with lipid and protein composition during de-etiolation of Arabidopsis seedlings.

## Results

### The photosynthetic machinery is functional after 14 h of de-etiolation

We analysed etioplast–chloroplast transition in Arabidopsis seedlings grown in the absence of exogenous sucrose for 3 days in darkness and then exposed to constant white light (Figure 1A). These experimental conditions were chosen to avoid effects of exogenous sucrose on seedling development and variations due to circadian rhythm. Upon illumination, the etiolated seedlings switched from the skotomorphogenic to the photomorphogenic developmental program, evidenced by opening of the apical hook and cotyledon greening and expansion (Figure 1B). We stopped the analysis following 96 h of illumination (T96), before the emergence of the primary leaves. Samples were collected at different selected time points during de-etiolation(Figure 1A).

**Figure 1:**
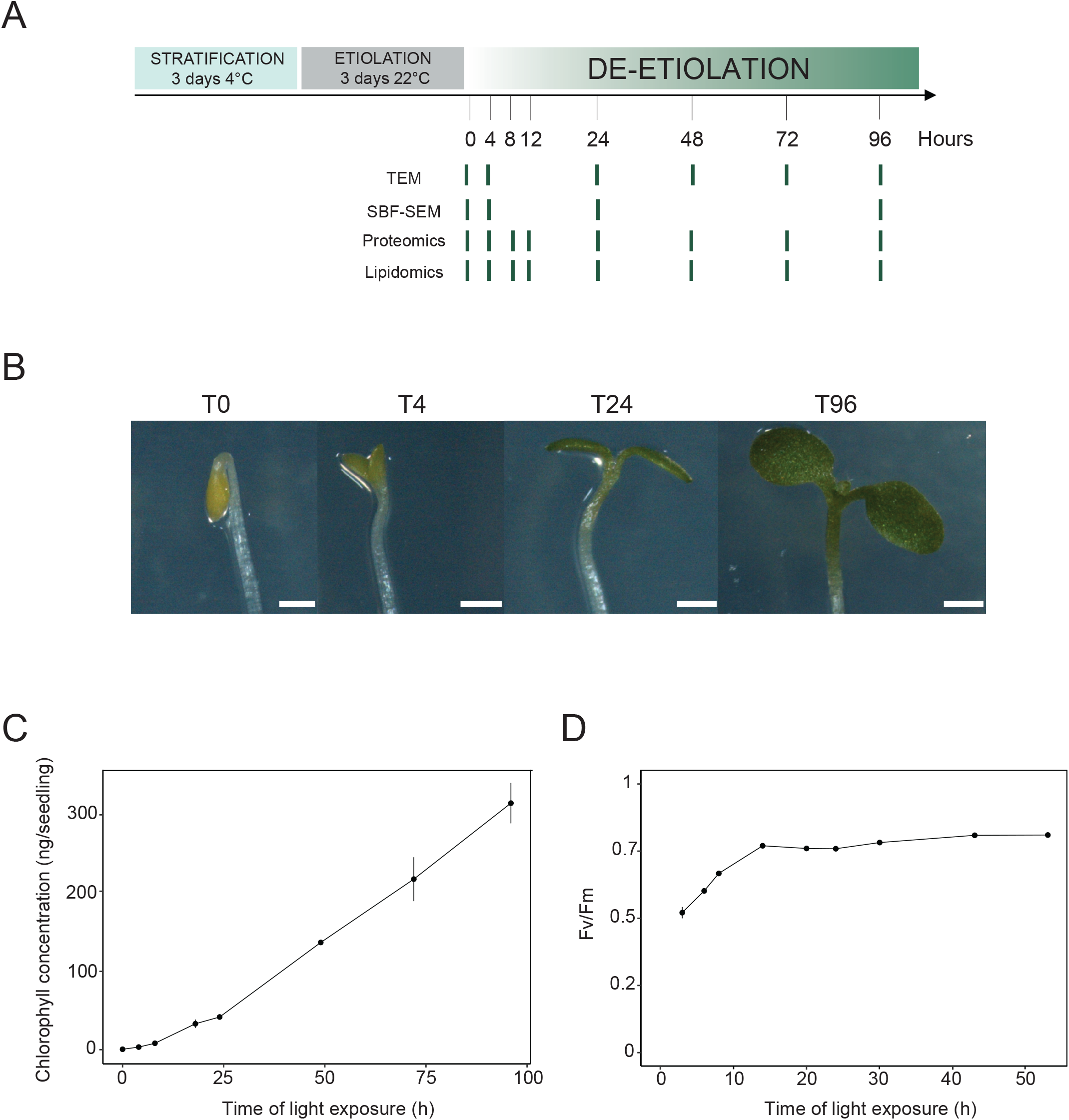
Photosynthesis onset during de-etiolation. (A) Scheme of the experimental design. Seeds of *Arabidopsis thaliana* (Columbia) sown on agar plates were stratified for three days at 4°C and then transferred to 22°C in the dark. After three days, etiolated seedlings were exposed to continuous white light (40 μmol/m^2^/s) and harvested at different time points during de-etiolation. Selected time points used for different analyses are indicated. (B) Cotyledon phenotype of etiolated seedlings (T0) after 4 h (T4), 24 h (T24), and 96 (T96) h in continuous white light. Scale bars: 0.5 mm. (C) Chlorophyll quantification at different time points upon illumination. Error bars indicate ± SD (n=3). (D) Maximum quantum yield of photosystem II (Fv/Fm). Error bars indicate ± SD (n=4–10). For some data points, the error bars are inferior to the size of the symbol. Measurements of further photosynthetic parameters are presented in Figure 1- figure supplement 1.

In angiosperms, chlorophyll synthesis arrests in the dark but starts immediately upon seedling irradiation (Von Wettstein et al., 1995). Chlorophyll levels in whole seedlings increased within the first 4 h of illumination (T4) and continued to increase linearly during subsequent illumination as the seedlings grew (Figure 1C). To evaluate photosynthetic efficiency during de-etiolation, we measured chlorophyll fluorescence and calculated the maximum quantum yield of PSII (Fv/Fm, Figure 1D and Figure 1- figure supplement 1). PSII maximum quantum yield increased during the initial period of illumination and was near the maximal value of 0.8 at 14 h of light exposure (T14), independent of light intensity (Figure 1D and Figure 1- figure supplement 1A). Other photosynthetic parameters (photochemical quenching, qP and PSII quantum yield in the light, ΦPSII, Figure 1-figure supplement 1 B and C) reached maximum values at T14 and remained stable thereafter, indicating that the assembly of fully functional photosynthetic machinery occurs within the first 14 h of de-etiolation, and that further biosynthesis of photosynthesis related compounds is efficiently coordinated.

### Major thylakoid structural changes occur within 24 h of de-etiolation

We determined the dynamics of thylakoid biogenesis during the etioplast–chloroplast transition by observing chloroplast ultrastructure in cotyledons using transmission electron microscopy (TEM) (Figure 2). Plastids present in cotyledons of etiolated seedlings displayed the typical etioplast ultrastructure with a paracrystalline PLB and tubular PTs (Figure 2A). The observed PLBs were constituted of hexagonal units with diameters of 0.8–1 μm (Figure 2E). By T4, the highly structured PLBs progressively disappeared and thylakoid lamellae were formed (Figure 2B). The lamellae were blurry and their thickness varied between 15 and 70 nm (Figure 2F). After 24 h of illumination (T24), the density of lamellae per chloroplast was higher than that at T4 due to an increase in lamellar length and number. Appressed regions corresponding to developing grana stacks also appeared by T24 (Figure 2C and G). These early grana stacks consisted of 2–6 lamellae with a thickness of 13 nm each (Figure 2- figure supplement 1). In addition, starch granules were present at T24, supporting the notion that these chloroplasts are photosynthetically functional and able to assimilate carbon dioxide (CO_2_). At T96, thylakoid membrane organisation was visually similar to that at T24, but with more layers per grana (up to 10 lamellae per grana; Figures 3D and H). In addition, singular lamella thickness at T96 increased by 2–3 nm compared to that at T24 (Figure 2- figure supplement 1). The major differences observed between T24 and T96 were increases in starch granule size and number and overall chloroplast size. Etioplast average length (estimated by measuring the maximum distance on individual slices) was 2 μm (± 0.9, n=10) in the dark (T0), whereas chloroplast average length was 6 μm (± 1.62, n=10) at T96 (Table 1). Collectively, these data show that photosynthetically functional thylakoid membranes form rapidly during the first 24 h of de-etiolation. This implies that there are efficient mechanisms for thylakoid assembly and structural organisation. Subsequent changes seem to involve the expansion of pre-existing structures (i.e. lamellae length and grana size) and the initiation of photosynthetic carbon fixation (reflected by starch content).

**Figure 2:**
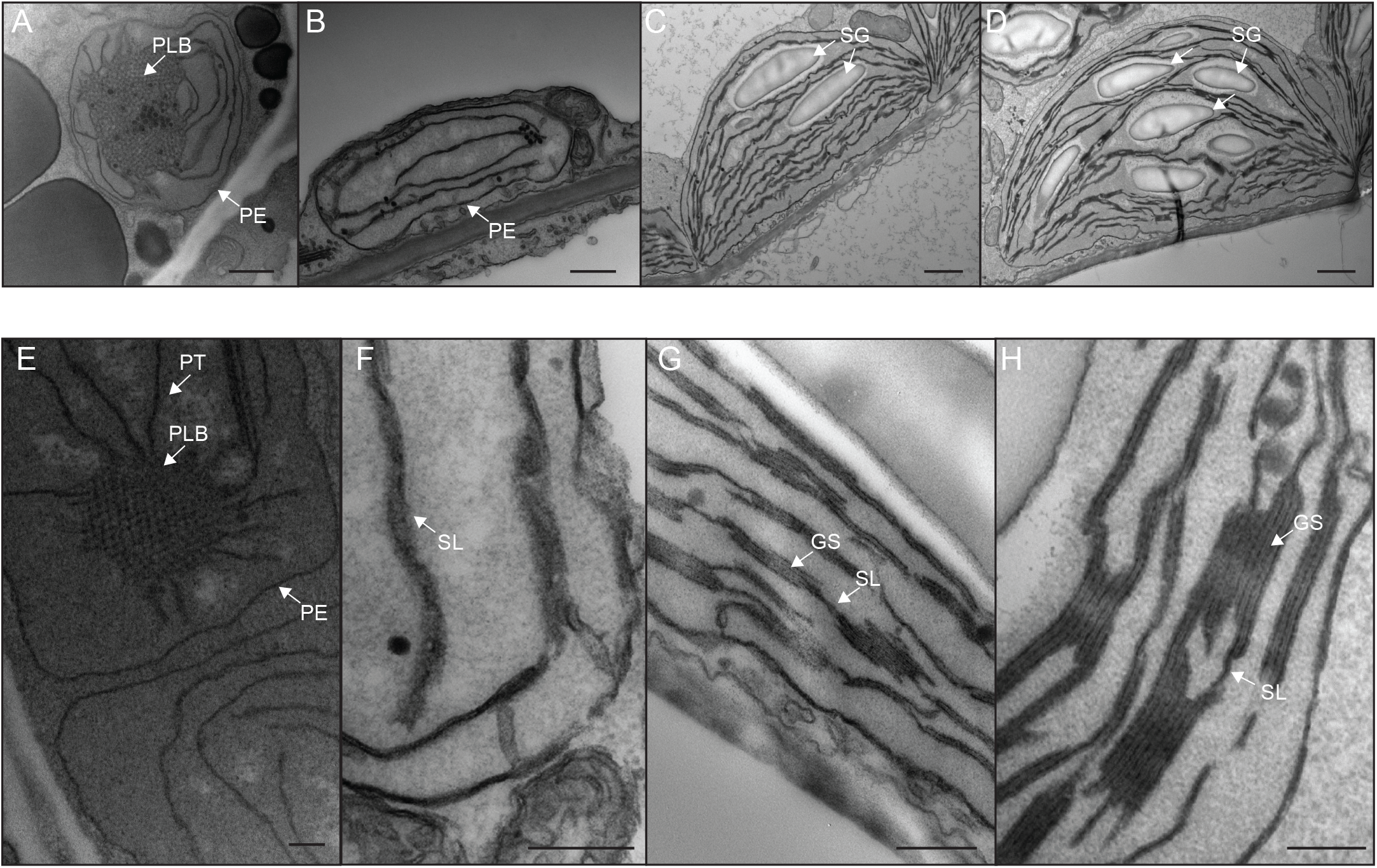
Qualitative analysis of chloroplast ultrastructure during de-etiolation. Transmission electron microscopy (TEM) images of cotyledon cells of 3-day-old, dark-grown *Arabidopsis thaliana* (Columbia) seedlings illuminated for 0 h (A and E), 4 h (B and F), 24 h (C and G), and 96 h (D and H) in continuous white light (40 μmol/m^2^/s). (A–D) Scale bars: 500 nm, (E–H) higher magnification of A–D images; Scale bars: 200 nm. PLB: prolamellar body; PT: prothylakoid; PE: plastid envelope; SG: starch grain; GS: grana stack; SL: single lamella. Specific details for measurements of lamella thickness are provided in Figure 2- figure supplement 1.

**Figure 3:**
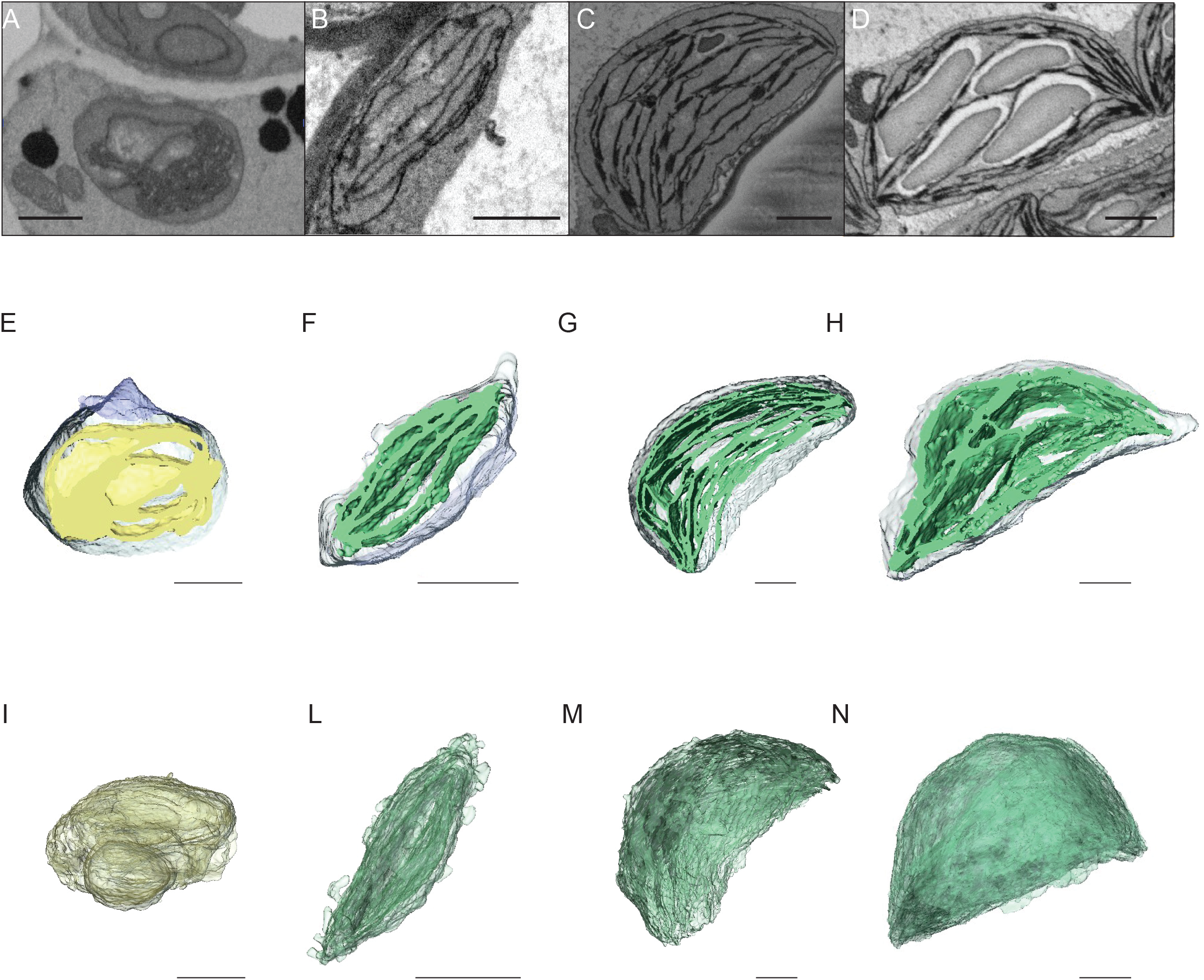
3D reconstructions of chloroplast thylakoid network during de-etiolation. (A–D) Scanning electron microscopy (SEM) micrographs of representative etioplasts and chloroplasts from 3-day-old, dark-grown *Arabidopsis thaliana* seedlings illuminated for 0 h (T0; A), 4 h (T4; B), 24 h (T24; C), and 96 h (T96; D) in continuous white light (40 μmol/m^2^/s). (E–H) Partial 3D reconstruction of thylakoid membranes (green) and envelope (blue) at T0 (E), T4 (F), T24 (G) and T96 (H). Z-depth of thylakoid membrane reconstruction corresponds to 0.06 μm (E), 0.10 μm (F), 0.13 μm (G), and 0.15 μm (H). (I–N). 3D reconstruction of a thylakoid membrane of an etioplast at T0 (I) or a chloroplast at T4 (L), T24 (M), and T96 (N). Scale bars = 1 μm. Details of grana segmentation at T24 are provided in Figure 3- figure supplement 1.

**Table 1.** Collection of quantitative data. Morphometric data corresponding to thylakoid surfaces and volumes, thylakoid/envelope surface ratio, and chloroplast and cell volumes were collected after 3View analysis. Chloroplast and cell volumes were also quantified by subsequent confocal microscopy analysis, whereas plastid length was measured using TEM images. Molecular data for galactolipids (GLs) were analysed by lipidomics, whereas PsbA, PsaC, and PetC were quantified by quantitative immunodetection.

### Quantitative analysis of thylakoid surface area per chloroplast during de-etiolation

To visualize entire chloroplasts and thylakoid networks in 3D, and to obtain a quantitative view of the total thylakoid surface area during chloroplast development, we prepared and imaged cotyledons at different developmental stages by SBF-SEM (Figure 3 A-D). PLBs, thylakoids, and envelope membranes were selected, and segmented images were used for 3D reconstruction (Figure 3E–N, and videos 1–4; see also Figure 2- figure supplement 1 and Figure 4- figure supplement 1 for grana segmentation). Similar to that observed by TEM (Figure 2), a drastic switch from PLB to thylakoid membrane occurred by T4: the typical structure of the PLB connected to PTs disappeared leaving only elongated lamellar structures (Figure 3E–F and videos 1 and 2). At T24 and T96, thylakoid membranes were organised in appressed and non-appressed regions and large spaces occupied by starch granules were observed (Figure 3G–H and videos 3 and 4). 3D reconstruction revealed a change in plastid shape from ovoid at T0 and T4 to hemispheric at T24 and T96 (Figure 3I–N).

Using 3D reconstruction of the thylakoid network for 3 or 4 chloroplasts for each developmental stage, quantitative data such as chloroplast volume and membrane surface area were extracted and calculated (Figure 4A and B, Figure 4 figure supplement 1 and Table 1). The total chloroplast volume increased about 11-fold from T4 (9.4 μm^3^) to T96 (112.14 μm^3^) (Table 1). In parallel, the thylakoid surface area increased about 30-fold reaching 2,086 (± 393) μm^2^ per chloroplast at T96 (Figure 4A and Table 1). The surface area increased drastically between T4 and T24 (about 22-fold) and much less (about 1.4-fold) between T24 and T96. Accordingly, quantification of the envelope surface area indicated that the ratio of the thylakoid to envelope surface area increased drastically from T4 to T24, but decreased slightly between T24 and T96 (Table 1).

**Figure 4:**
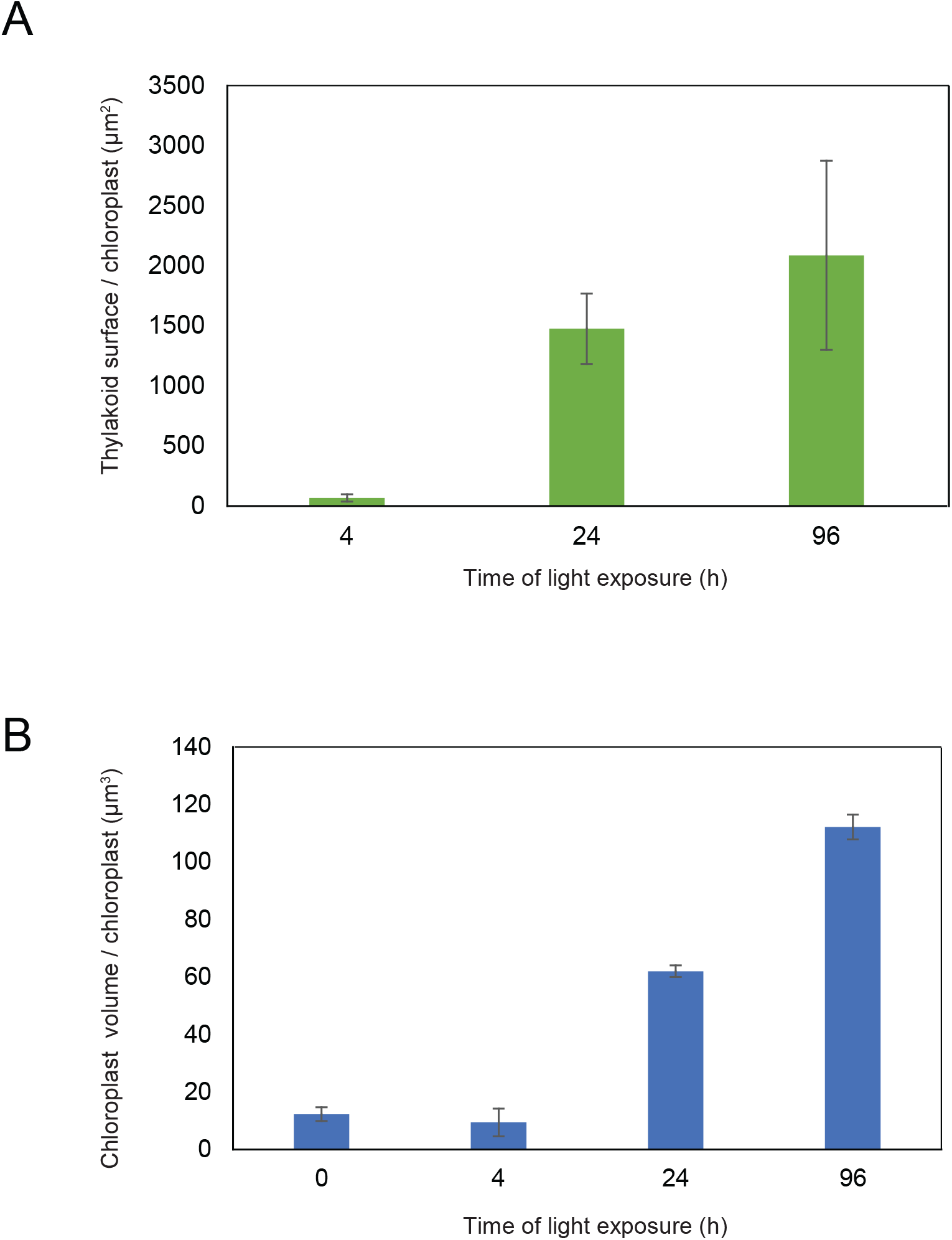
Quantitative analysis of chloroplast volume and thylakoid surface during de-etiolation. Quantification of thylakoid surface per chloroplast (A) and chloroplast volume (B) using 3-day-old, dark-grown *Arabidopsis thaliana* (Columbia) seedlings illuminated for 0 h, 4 h, 24 h, and 96 h in continuous white light (40 μmol/m^2^/s). Morphometric data were quantified by Labels analysis module of Amira software. Error bars indicate ± SD (n=3). The total thylakoid surface indicated in A corresponds to the thylakoid surface exposed to the stroma, calculated in Amira software, in addition to the percentage of the grana surface (%Gs) calculated as described in Figure 3- figure supplement 1.

Our observations indicated that chloroplast development during the first 96 hours of de-etiolation could be separated into two phases: a first phase reflected by qualitative changes (i.e. structure establishment and reorganisation of the thylakoid network architecture) and a second phase (starting before T24) during which thylakoid surface increased due to the expansion and stacking of lamellae. We further analysed these temporal processes at the molecular level focusing on proteins and lipids that constitute the thylakoid membrane.

### Dynamics of plastid proteins related to thylakoid biogenesis

We analysed the full proteome to reveal the dynamics of protein accumulation during de-etiolation. Total proteins were prepared from 3-day-old etiolated seedlings exposed to light for 0–96 h (eight time points; Figure 1A) and quantified by label-free shot-gun mass spectrometry. For relative quantification of protein abundances between different samples, peptide ion abundances were normalized to total protein (see Materials and Methods). We considered further only those proteins that were identified with a minimum of two different peptides (with at least one being unique; see Methods for information on protein grouping), resulting in the robust identification and quantification of more than 5,000 proteins. Fold changes of protein abundances between two time points were regarded as significant if their adjusted *p*-value (i.e. the *q*-value) was < 0.01.

The first 12 h of illumination (T12) saw very few significant changes in protein abundance (Supplemental Dataset 1). After 8 h of illumination (T8), we observed decreased abundance of only one protein (the photoreceptor cryptochrome 2, consistent with its photolabile property) and increased levels of only three proteins, which belonged to the chlorophyll a/b binding proteins category involved in photoprotection (AT1G44575 = PsbS; AT4G10340= Lhcb5; AT1G15820= Lhcb6; (Chen et al., 2018; Li et al., 2000). A drastic change of proteome composition occurred by T24, with 467 proteins showing a significant increase in abundance with over 2-fold change (FC>2) compared with the etiolated stage, and 150 proteins showing a significant decrease with over 2-fold change (FC<0.5). As expected, the 100 most-upregulated proteins comprised proteins related to photosynthesis, proteins constituting the core and antennae of photosystems, and proteins involved in carbon fixation (Supplemental Dataset 1).

To monitor the dynamics of the plastidial proteome, we selected proteins predicted to localize to the plastid (consensus localization from SUBA4; Hooper et al., 2017). Generation of a global heatmap for each of the 1,112 potential plastidial proteins revealed different accumulation patterns (Supplemental Dataset 2 and Figure 5- figure supplement 1). Hierarchical clustering showed a categorization into six main clusters. Cluster 1 (purple) contained proteins whose relative amounts decreased during de-etiolation. Clusters 2, 5, and 6 (pink, light green, and dark green, respectively) contained proteins whose relative amounts increased during de-etiolation but differed with respect to the amplitude of variations. Proteins in clusters 2 and 6 displayed the largest amplitude of differential accumulation. Gene ontology (GO) analysis (Mi et al., 2019) indicated a statistically significant overrepresentation of proteins related to the light reactions of photosynthesis in clusters 2 and 6 (Supplemental Dataset 2). Underrepresentation of organic acid metabolism, in particular carboxylic acid metabolism, characterized cluster 2, whereas overrepresentation of carboxylic acid biosynthesis and underrepresentation of photosynthetic light reactions were clear features of cluster 3. Protein levels in cluster 3 changed only moderately during de-etiolation in contrast with proteins levels in cluster 2. No biological processes were significantly over- or underrepresented in clusters 1, 4, and 5.

To analyse the dynamics of proteins related to thylakoid biogenesis, we selected specific proteins and represented their pattern of accumulation during de-etiolation (Figure 5). We included proteins constituting protein complexes located in thylakoids (complexes constituting the electron transport chain and the ATP synthase complex) and proteins involved in chloroplast lipid metabolism, chlorophyll synthesis, and protein import into the chloroplast. In agreement with that depicted in the global heatmap (Figure 5- figure supplement 1), all photosynthesis-related proteins increased in abundance during de-etiolation (Figure 5A). However, our hierarchical clustering did not show any particular clustering per complex. Only few chloroplast-localized proteins related to lipid biosynthesis were present in our proteomics data set. Among the eight detected proteins, two appeared differentially regulated; fatty acid binding protein 1 (FAB1) and fatty acid desaturase 7 (FAD7) levels increased only between 72 h of illumination (T72) and T96, whereas the other proteins gradually accumulated over the course of de-etiolation (Figure 5B). Etioplasts initiate synthesis of chlorophyll precursors that are blocked at the level of protochlorophyllide synthesis, with protochlorophyllide oxidoreductase A (PORA) in its inactive form accumulating to high levels in the etioplast before subsequently decreasing at the protein level upon activation and degradation following light exposure (Blomqvist et al., 2008; Runge et al., 1996; Von Wettstein et al., 1995). In agreement, illumination resulted in increased amounts of all detected proteins of the chlorophyll biosynthesis pathway, except PORA, which clearly decreased and was separated from other chlorophyll-related proteins (Figure 5C). We also selected proteins involved in protein import in chloroplasts, focusing on the TOC-TIC machinery (Figure 5D) that is the major route for plastid protein import and essential for chloroplast biogenesis (Kessler and Schnell, 2006). Past studies identified several TOC preprotein receptors that are proposed to display differential specificities for preprotein classes (Bauer et al., 2000; Bischof et al., 2011). The composition of plastid import complexes varies with developmental stages and in different tissues, thereby adjusting the selectivity of the import apparatus to the demands of the plastid and influencing its proteome composition (Demarsy et al., 2014; Kubis et al., 2003). Accordingly, the TOC receptors TOC120 and TOC132, which are important for the import of proteins in non-photosynthetic tissues, were more abundant in etioplasts compared to fully-developed chloroplasts (compare T0 and T96). TOC120 and TOC132 were part of a cluster separated from other components of the plastid machinery, such as the TOC159 receptor associated with large-scale import of proteins in chloroplasts. The general import channel TOC75 (TOC75 III) maintained stable expression levels throughout de-etiolation, reflecting its general role in protein import. All other components clustered with TOC159 and displayed gradual increases in accumulation during de-etiolation. Most of these components have not been reported to confer selectivity to the import machinery, which suggests an overall increase of chloroplast protein import capacity.

**Figure 5:**
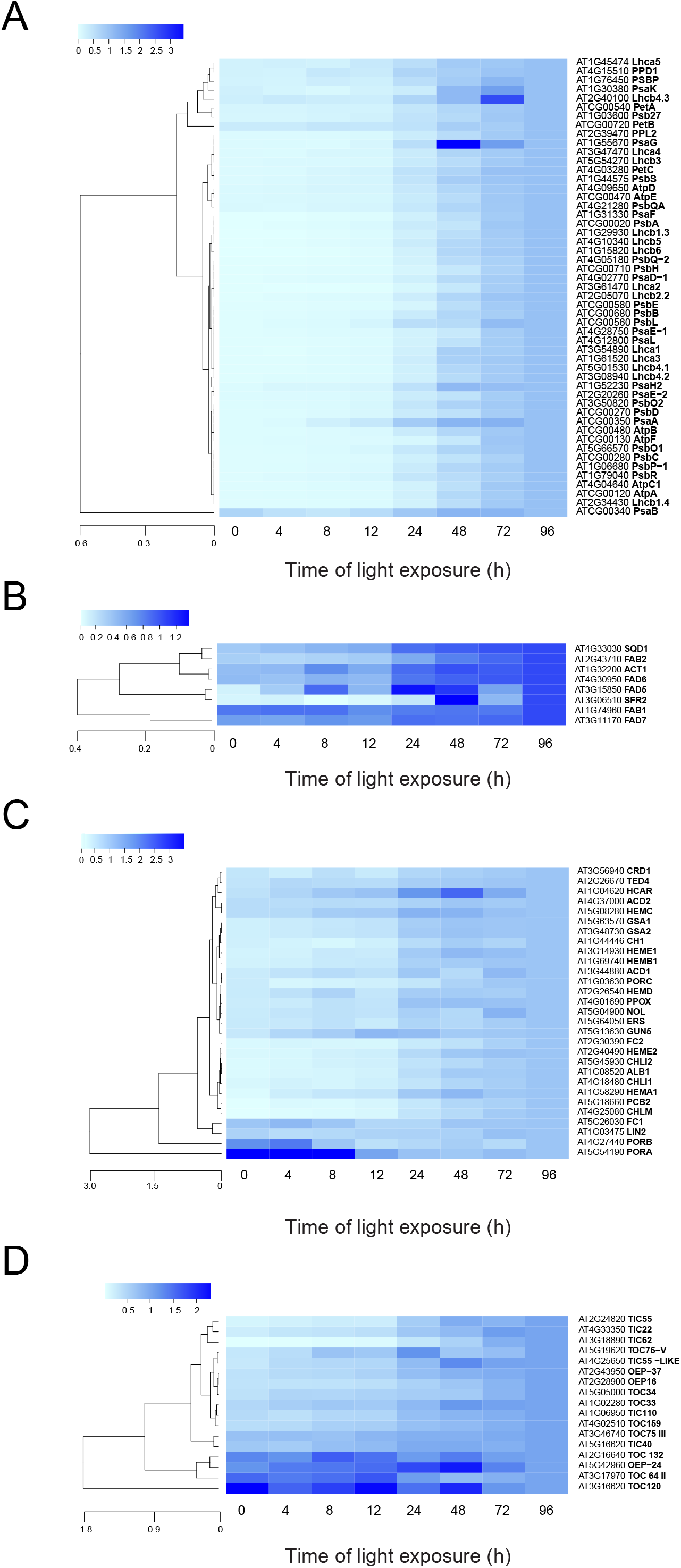
Accumulation dynamics of plastid proteins during de-etiolation. 3-day-old etiolated seedlings of *Arabidopsis thaliana* were illuminated for 0 h (T0), 4 h (T4), 8 h (T8), 12 h (T12), 24 h (T24), 48 h (T48), 72 h (T72), and 96 h (T96) under white light (40 μmol/m^2^/s). Hierarchical clustering (Euclidean, average linkage) of normalized protein abundance for photosynthesis-(A), galactolipid metabolism-(B), chlorophyll metabolism-(C), and protein import-related proteins during de-etiolation (D). Protein abundance was quantified by shot-gun proteomics and heatmap colors indicate the fold change (average of 3–4 replicates) of each selected protein at each time point of de-etiolation (T0 to T96), relative to the last time point (T96). Note that some PORA values in panel D were higher than 3.5 and outside of the color range limits. Further hierarchical clustering based on the accumulation dynamics of all plastid-localized proteins is provided in Figure 5- figure supplement 1.

To validate and complement our proteomic data, we used immunoblot analysis to detect and quantify representative proteins of the photosynthetic complexes. Overall, immunoblot and proteomics provided similar results (Figure 6 and Figure 6- figure supplement 1). PsbA and PsbD (PSII reaction center core), PsbO (Oxygen Evolving Complex), and Lhcb2 (outer antenna complex) proteins were detectable in seedlings at T0, gradually increasing thereafter. Accumulation of the PSI proteins PsaC and PsaD and the Cyt *b*_6_*f* complex protein PetC started later; these proteins were detectable starting at T8 (Figure 6A and Figure 6- figure supplement 1). Interestingly, AtpC (ATP synthase complex) was detectable in the etioplast, as described previously (Plöscher et al., 2011). Other proteins were selected as markers of etioplast– chloroplast transition. As expected, ELIPs (Early Light Induced Protein) transiently accumulated upon the dark-to-light transition (Figure 6A) (Kimura et al., 2003). As in the proteome analysis, PORA accumulated in etiolated seedlings (T0) and then progressively disappeared upon light exposure. We performed absolute quantification for PsbA, PsaC, and PetC proteins using recombinant proteins as standards (Figure 6B and C and Figure 6- figure supplement 1). Quantitative data (nmol/seedling) were obtained and normalized using the last time point (Figure 6C) to compare the dynamics of protein accumulation. In addition, the comparison of PsbA and PsaC (representative proteins of PSII and PSI, respectively) showed that PsbA levels were about twice that of PsaC at T96 (Figure 6B and C).

**Figure 6:**
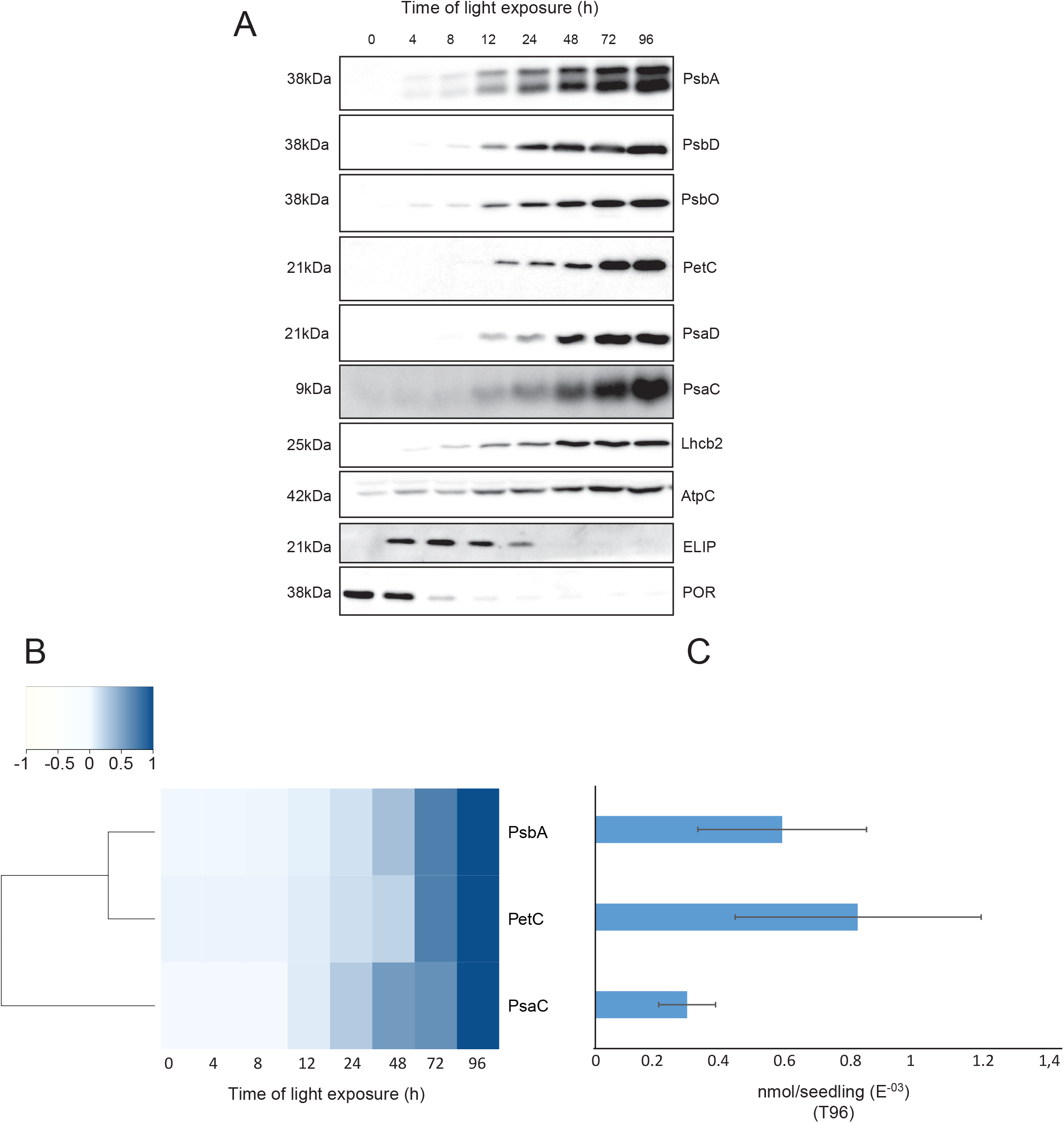
Accumulation dynamics of photosynthesis-related proteins during de-etiolation. 3-day-old etiolated seedlings of *Arabidopsis thaliana* were illuminated for 0 h (T0), 4 h (T4), 8 h (T8), 12 h (T12), 24 h (T24), 48 h (T48), 72 h (T72), and 96 h (T96) under white light (40 μmol/m^2^/s). (A) Proteins were separated by SDS-PAGE and transferred onto nitrocellulose membrane and immunodetected with antibodies against PsbA, PsbD, PsbO, PetC, PsaD, PsaC, Lhcb2, AtpC, ELIP, POR proteins. (B–C) Quantification of PsbA, PetC, and PsaC during de-etiolation. Heatmap (B) was generated after normalization of the amount of each protein relative to the last time point (T96). Graph (C) corresponds to the absolute quantification of proteins at T96. Error bars indicate ± SD (n=3). Quantification of photosystem-related proteins during de-etiolation is detailed in Figure 6- figure supplement 1.

### Dynamics of chloroplast membrane lipids

Total lipids were extracted from seedlings collected at different time points during de-etiolation (T0, T4, T8, T12, T24, T48, T72, and T96), analysed by ultra-high pressure liquid chromatography–mass spectrometry (UHPLC-MS), and quantified against pure standards (supplemental Dataset 3). We analysed the quantity and kinetics of accumulation of 12 different species of galactolipids (Figure 7A and B). MGDG 18:3/16:3, MGDG 18:3/18:3, MGDG 18:3/16:1, DGDG 18:3/18:3, and DGDG 18:3/16:0 were the most abundant lipids detected at all time points. Accumulation of all galactolipids increased upon de-etiolation; however, clustering analysis identified two distinct kinetic patterns. One group displayed a leap between T8 and T12, whereas the other group showed a more gradual increase during the de-etiolation period (Figure 7C). Interestingly, the two clusters separated the lipids according to the two pathways described for galactolipid synthesis, namely the ER and PL pathways (Figure 7A and B) (Marechal et al., 1997; Ohlrogge and Browse, 1995). During early stages of de-etiolation (T0–T24), we observed an incremental accumulation of MGDG and DGDG galactolipids derived from the ER pathway, whereas galactolipids from the PL pathway started to accumulate at T24 (Figure 7A and B). The MGDG/DGDG ratio decreased between T0 and T8. This was associated with the transition from PLB (cubic lipid phase) to thylakoid membrane (lamellar structure) (Bottier et al., 2007). The MGDG/DGDG ratio started to increase gradually at T8 and was constant by T72 and T96 (Figure 7D).

**Figure 7:**
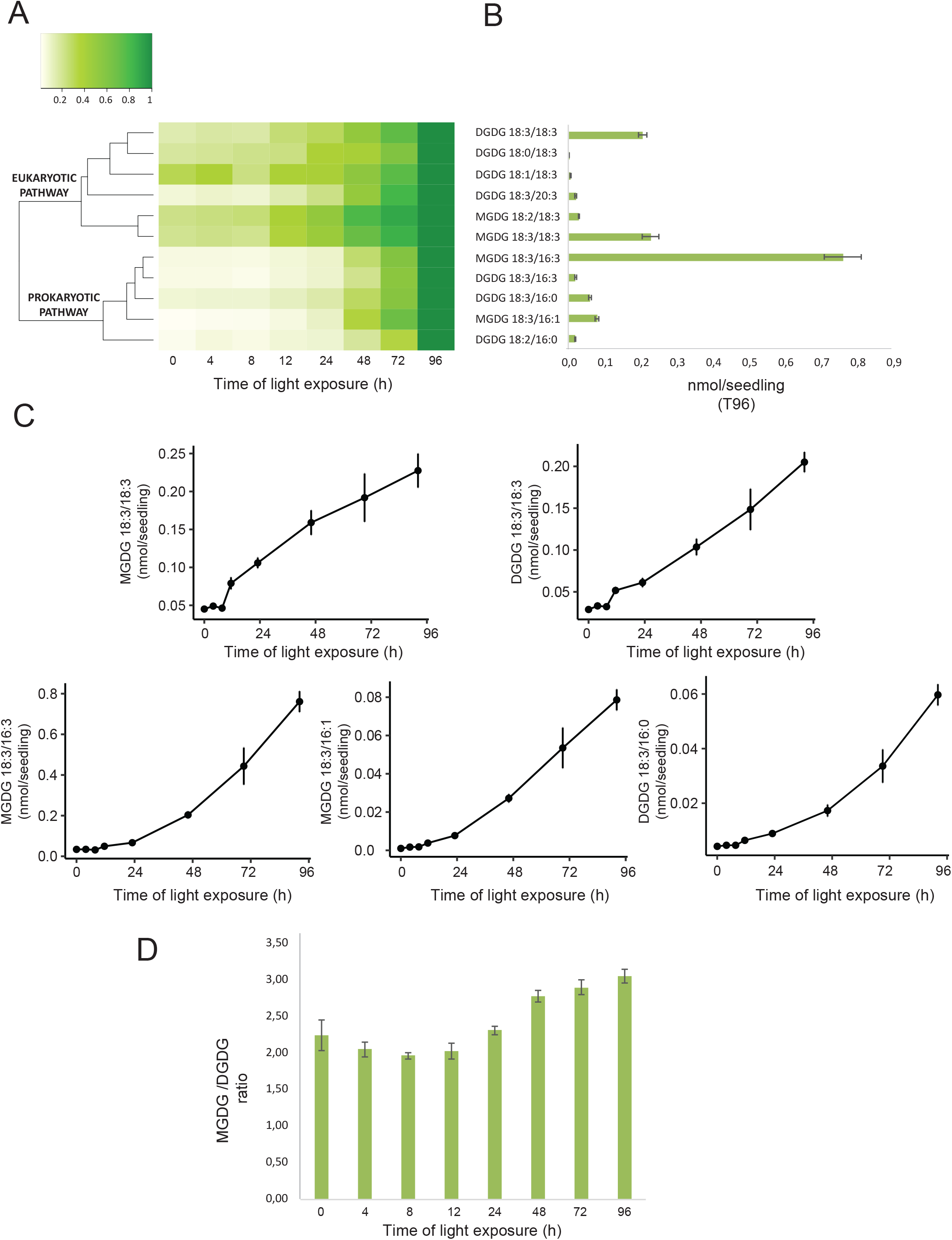
Accumulation dynamics of galactolipids during de-etiolation. 3-day-old etiolated seedlings of *Arabidopsis thaliana* were illuminated for 0 h (T0), 4 h (T4), 8 h (T8), 12 h (T12), 24 h (T24), 48 h (T48), 72 h (T72), and 96 h (T96) under white light (40 μmol/m^2^/s). (A) Heatmap representation of galactolipids (MGDG and DGDG) during de-etiolation. Samples were normalized to the last time point (T96). (B) Absolute quantification at T96 expressed in nmol/seedling. Error bars indicate ± SD (n=4). (C) Absolute quantification (nmol/seedling) of the most abundant chloroplast galactolipids MGDG (MGDG 18:3/18:3, MGDG 18:3/16:3, MGDG 18:3/16:1) and DGDG (DGDG 18:3/18:3, DGDG 18:3/16:0) at different time points during de-etiolation. Error bars indicate ± SD (n=4). (D) The MGDG/DGDG ratio was calculated using all 12 species of galactolipids detected during de-etiolation. Error bars indicate ± SD (n=4).

### Identification of a chloroplast division phase

We observed a massive increase in the accumulation of photosynthesis-related proteins and galactolipids between T24 and T96, corresponding to FC>2 in the levels of all major chloroplast proteins and lipids (Figures 6 and 7). Intriguingly, the total thylakoid surface per chloroplast increased by only 41 % between these two time points (Figure 4A and Table 1). We reasoned that the increase in chloroplast proteins and lipids between T24 and T96 could be explained by increased chloroplast number (per cell and thus per seedling) and thus total thylakoid surface per seedling. We therefore determined chloroplast number per cell and the cell number and volume for each developmental stage through SBF-SEM analysis (T0, T4, T24, and T96) and confocal microscopy analysis for intermediary time points (T24–T96) (Figure 8 and Figure 8- figure supplement 1). The chloroplast number per cell was constant from T4 (25 ± 8) to T24 (26 ± 6); however, in parallel with cell expansion (Figure 8A and B), chloroplast number increased sharply (4-fold increase) between T24 (26 ± 6) and T96 (112 ± 29), indicating that two rounds of chloroplast division occurred during this time. Immunoblot analysis of FILAMENTOUS TEMPERATURE-SENSITIVE FtsZ1, FtsZ2-1, and FtsZ2-2 proteins showed that these key components of the chloroplast division machinery were already present during the early time points of de-etiolation. We observed considerably increased accumulation of these proteins between T24 and T48, consistent with the idea that activation of chloroplast division takes place at T24, leading the proliferation of chloroplasts (Figure 8C–D). However, levels of ACCUMULATION AND REPLICATION OF CHLOROPLAST 5 (ARC5) protein, another key component of the chloroplast division machinery, clearly increased during de-etiolation between T8 and T12, presumably reflecting assembly of the chloroplast division machinery before its activation and the proliferation of chloroplasts (Figure 8D). To test whether there is a correlation between chloroplast division and either volume or developmental stage, we measured the volume of dividing chloroplasts at T24 and T96 using images acquired by SBF-SEM (Figure 8E and Figure 4B). The average size of dividing chloroplasts at T24 was higher than the average size of all chloroplasts (96 μm^3^ compared to 62 μm^3^). The volume of dividing chloroplasts at T96 was consistently higher than 100 μm^3^ although some of the chloroplasts present were smaller (Figure 8E and Figure 4B). Altogether, this indicates that developing chloroplasts only divide once a certain chloroplast volume is reached.

**Figure 8:**
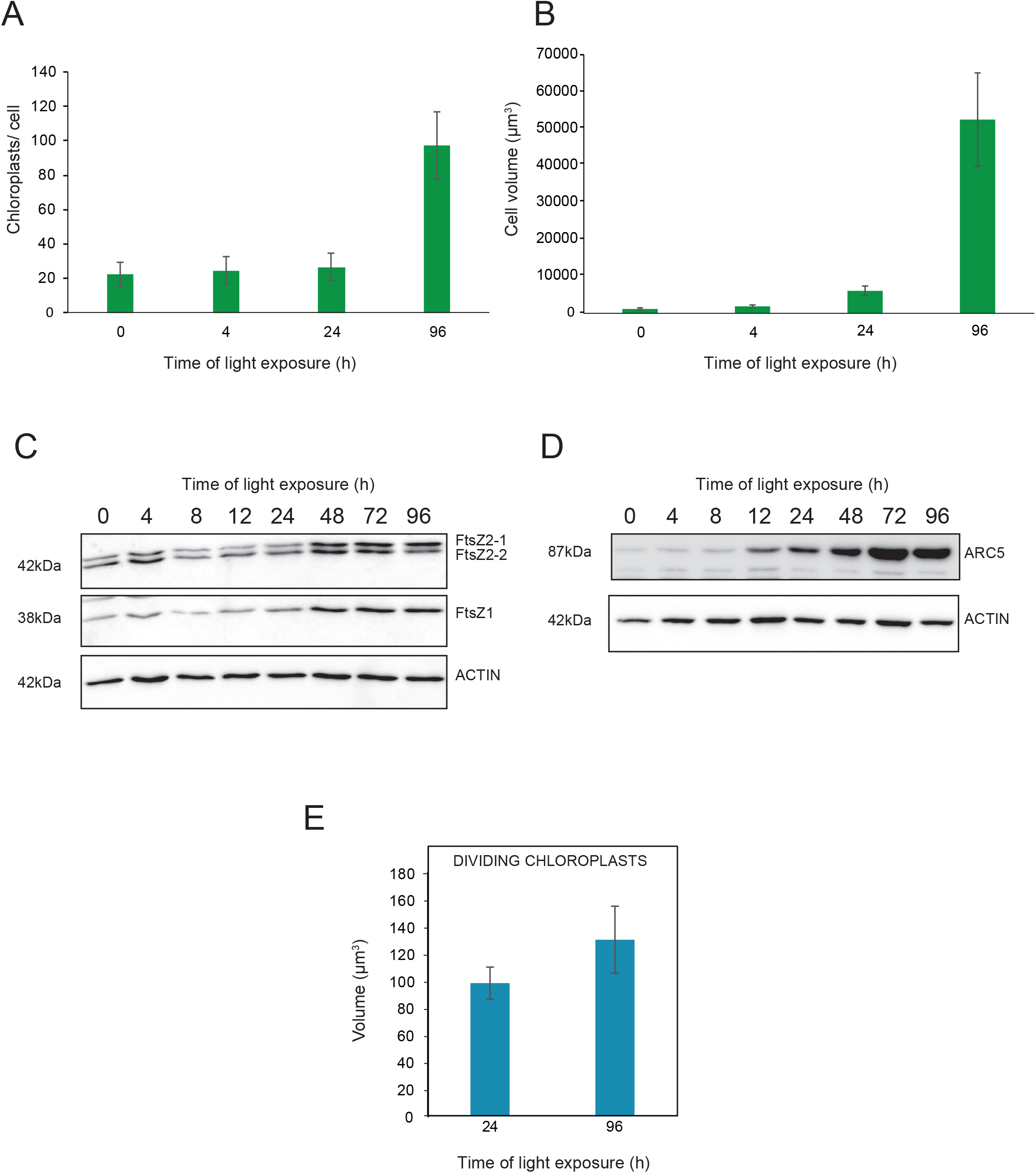
Relationship between chloroplast proliferation and chloroplast volume. (A-B) Chloroplast number and cell volume in cotyledons of 3-day-old, dark-grown *Arabidopsis thaliana* seedlings illuminated for 0 h (T0), 4 h (T4), 24 h (T24), and 96 h (T96) in continuous white light (40 μmol/m^2^/s). (A) Chloroplast number per cell during de-etiolation. Error bars indicate ± SD (n=6 for T0 and T7; 7 for T24; 5 for T96). (B) Cell volume was quantified by the Labels analysis module of Amira software. Error bars indicate ± SD (n=5–6). (C–D) Total proteins were extracted from T0–T96 seedlings, separated on SDS-PAGE, and transferred onto nitrocellulose. Proteins involved in plastid division (C, FtsZ; D, ARC5) and loading control (actin) were detected using specific antibodies (FtsZ2 antibody recognizes both FtsZ2-1 and FtsZ2-2). (E) Volume of dividing chloroplast at T24 and T96. Error bars indicate ± SD (n=3). Further details of chloroplast proliferation in parallel with cell expansion are provided in Figure 8- figure supplement 1.

### Model of thylakoid surface expansion over time

During de-etiolation, thylakoid surface increased with the accumulation of galactolipids and photosynthesis-related proteins, leading to the formation of functional chloroplasts. To determine the thylakoid membrane surface area per seedling and its expansion over time, we first calculated the surface area occupied by the main galactolipids (MGDG and DGDG) and photosynthetic complexes (PSII, Cyt *b*_6_*f* and PSI) per seedling (Table 2).

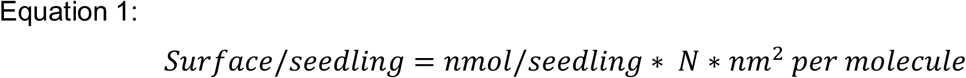

**Table 2.** Surface area occupied by the main galactolipids (MGDG and DGDG) and photosynthetic complexes (PSII, cyt*b*_6_*f*, and PSI). Shown are values at different time points following illumination of 3-day-old etiolated seedlings. Each value (in bold) indicates the calculated surface area in μm^2^ and corresponds to the average of three biological replicates. Errors indicate SD.

Quantitative data for MGDG, DGDG, PsbA, PetC, and PsaC (nmol/seedling) obtained from lipidomic and immunological analyses (Figures 6 and 7) were converted into number of molecules/seedling using the Avogadro constant (*N*). To calculate the surface area exposed to the stroma and account for the lipid double layer of the membrane, corresponding values of lipids (Figure 4A) were divided by 2. In addition, the lipid values were corrected by subtracting the portion of lipids incorporated into the envelope rather than present in the thylakoids (Table 1, Table 2 and Supplemental Dataset 3). The surface area occupied by molecules of MGDG and DGDG, and that of PSII, Cyt *b*_6_*f*, and PSI photosynthetic complexes (nm^2^ per molecule, corresponding to stroma-exposed surface) were retrieved from the literature (Table 3). Specifically, we used the minimal molecular area of MGDG and DGDG (Bottier et al., 2007). To quantify the surface area occupied by the galactolipids and photosynthetic complexes in thylakoids per seedling, the number of molecules per seedling of galactolipids was multiplied by the corresponding molecular surface area, whereas the number of molecules per seedling of PsbA, PetC, and PsaC (subunits of PSII, Cyt *b*_6_*f*, and PSI, respectively) were multiplied by the surface area of the corresponding complex (see Table 3).

**Table 3.** Surface area occupied by galactolipid and photosynthetic complexes. (A) Values were retrieved from the corresponding references. MGDG and DGDG surfaces correspond to the minimal molecular area. The surfaces of PSII-LHCII, PSI, and Cyt *b*_6_*f* complexes correspond to the surface exposed to the stroma (19*26 nm, 20*15 nm, and 90*55 Å, respectively). (B) Values from the table in panel A were used to calculate the total surface per seedling corresponding to MGDG and DGDG galactolipids, and PSII, PSI, and Cyt *b*_6_*f* complexes.

We calculated thylakoid surface (S) per seedling fir each time point (t) as the sum of the surface occupied by MGDG, DGDG, photosynthetic complexes (PS), and ɛ per seedling, the latter of which corresponds to compounds such as other lipids (e.g. sulfoquinovosyldiacylglycerol, plastoquinone) or protein complexes (ATP synthase and NDH) that were not quantified.

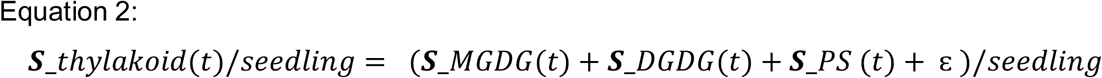

Omitting the unknown ɛ factor, we plotted the thylakoid surface calculated for each time point where quantitative molecular data were available (T0, T4, T8, T12, T24, T48, T72, and T96) as a function of the duration of light exposure (Figure 9- figure supplement 1). The best fitting curve corresponded to a S-shaped logistic function, characterized by a lag phase at early time points (T0–T8), followed by a phase of near-linear increase, and a final plateau at the final time points (T72–T96). To model this function, a four-parameter logistic non-linear regression equation was used to describe the dynamics of the total thylakoid surface over time (Figure 9- figure supplement 1C).

### Superimposition of molecular and morphometric data

We compared the values of thylakoid surface, as obtained with the model based on molecular data, with the values obtained from the morphometric analysis (Figure 9). The total thylakoid surface per seedling (S_thylakoid_morpho) was calculated by multiplying the thylakoid surface (S_thylakoid) per chloroplast obtained by morphometrics (Figure 4A) by the number of chloroplasts (nb.cp) per cell (Figure 8A) and the number of cells (nb.cells) per seedlings for each time point (t).

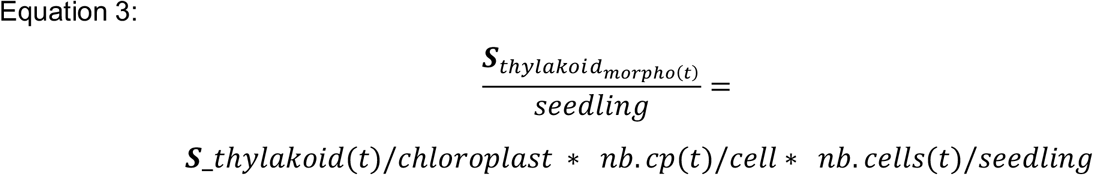

**Figure 9:**
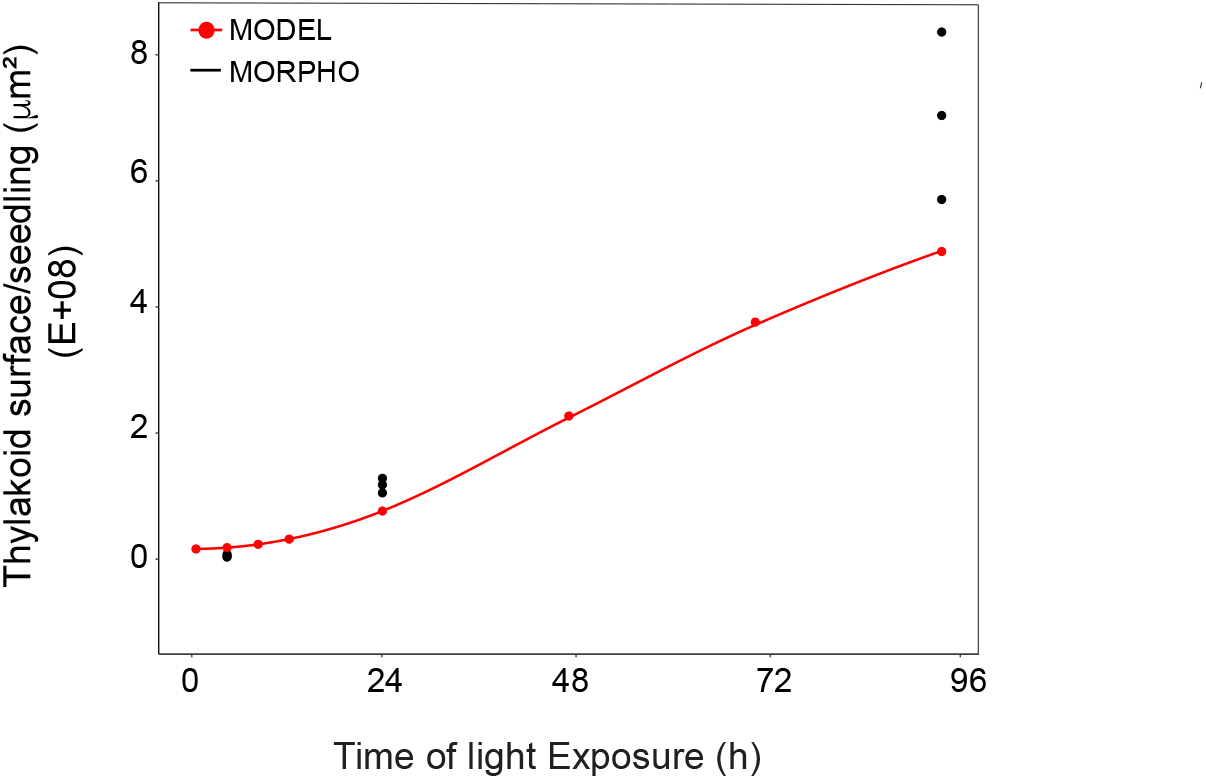
Superimposition of thylakoid surface per seedling obtained from morphometric analysis and mathematical modeling. Thylakoid surface per seedling was estimated using quantitative data from 3View analysis (‘MORPHO’ black dots at T4, T24, and T96; and see Figure 4 and Table 1) and model generated using the quantitative data from proteomics and lipidomics (‘MODEL’ red line at T0, T4, T8, T12, T24, T48, T72, and T96, and Table 1). Further details are provided in Figure 9- figure supplement 1 and 2.

We estimated cell number per seedling by measuring the total volume occupied by palisade and spongy cells in cotyledons (that corresponded to 50% of total cotyledon volume) (Figure 9- figure supplement 2) and dividing this by the average cell volume quantified by Amira software (Figure 7B). As reported previously (Pyke and Leech, 1994), cell number was constant during cotyledon development. We estimated this number as 3,000 mesophyll and palisade cells per seedling at T24 and T96 (Figure 9- figure supplement 2). The thylakoid membrane surface quantified by the morphometric approach was also estimated at T4, assuming that cell number per cotyledon remained similar between T4 and T24. We compared the thylakoid surface predicted by our mathematical model to the surface estimated experimentally with our 3D thylakoid reconstruction and morphometric measurements (Figure 9 and Table 1). As shown in Figure 9, the two approaches showed very similar total thylakoid surface area per seedling at T4 and T24 and differences in this parameter by T96.

## Discussion

Here, the analysis of 3D structures of entire chloroplasts in Arabidopsis in combination with proteomic and lipidomic analyses provide an overview of thylakoid biogenesis. Figure 10 depicts a summary of the changes that occur during the de-etiolation process. When considering chloroplast development, our study shows that de-etiolation is divided into two phases. We documented structural changes (disassembly of the PLB and the gradual formation of thylakoid lamellae) and initial increases of ER- and PL-pathway galactolipids and photosynthesis-related proteins (PSII, PSI, and Cyt *b*_6_*f*) during the ‘Structure Establishment Phase’, which was followed by increased chloroplast number in parallel with cell expansion in the ‘Chloroplast Proliferation Phase’. Collection of quantitative data allowed us to create a mathematical model of thylakoid membrane expansion and describe this process during de-etiolation.

**Figure 10:**
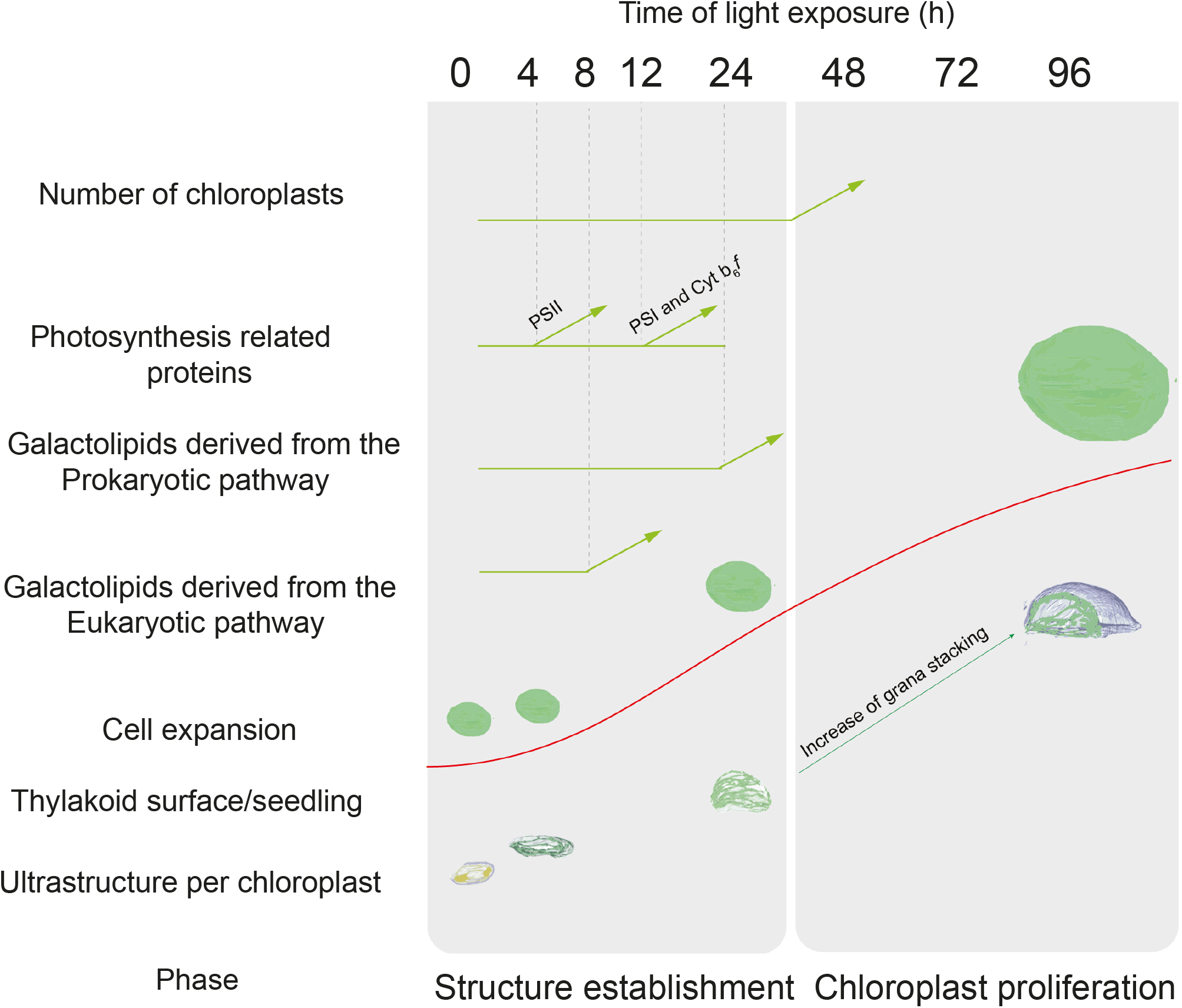
Overview of changes observed during the de-etiolation process in *Arabidopsis thaliana* seedlings. The ‘Structure Establishment Phase’ is correlated with disassembly of the PLB and gradual formation of the thylakoid membrane as well as an initial increase of eukaryotic (after 8 h) and prokaryotic (after 24 h) galactolipids and photosynthesis-related proteins (PSII subunits at 4 h, PSI and cyt b_6_*f* at 12 h). The subsequent ‘Chloroplast Proliferation Phase’ is associated with an increase in chloroplast number in concomitance with cell expansion, a linear increase of prokaryotic and eukaryotic galactolipids and photosynthesis-related proteins, and increased grana stacking. The red curve (retrieved from the Figure 9) shows thylakoid surface/seedling dynamics during the de-etiolation process.

### A set of 3D reconstructions of whole chloroplasts by SBF-SEM

In contrast to electron tomography, which is limited in the volume of observation, SBF-SEM allows the acquisition of ultrastructural data from large volumes of mesophyll tissue and the generation of 3D reconstructions of entire cells and chloroplasts (Figure 3 and Figure 8- figure supplement 1). SEM image resolution was sufficient to visualize stromal lamellae and grana contours, whereas grana segmentation in different lamellae was deduced according to our own TEM analysis and literature data (Figure 2- figure supplement 1 and Figure 3- figure supplement 1). This approach allowed us to obtain quantitative data of chloroplast and thylakoid structure at different developmental stages during de-etiolation at the whole-chloroplast level. By T96, the latest time point of our analysis, the total surface area of thylakoids present in the seedling cotyledons was about 700 mm^2^ (see values in Table 1 for calculation), about 500-fold greater than the surface area of one cotyledon at this developmental stage. This result is supported by previous estimates made regarding thylakoid surface area relative to leaf surface area (Bastien et al., 2016; Demé et al., 2014). Moreover, the extent of thylakoid surface area emphasizes how fast and efficient thylakoid biogenesis is during plant development, allowing plants to optimize light absorption capacity, ensuring their primary source of energy.

### Chloroplast development: ‘Structure Establishment Phase’

We observed TEM images and quantified 3D chloroplast ultrastructure by SBF-SEM analysis during chloroplast differentiation. Typical etioplast structure of the PLB connected with tubular PTs was replaced by lamellar thylakoids by T4. Measurements of PLB diameter and thylakoid length and thickness were comparable with literature values (Biswal et al., 2013; Daum et al., 2010; Kirchhoff et al., 2011), indicating that these morphometric values are conserved between various model organisms. Thylakoid surface increased 20-fold between T4 and T24. Remarkably, PSII maximum quantum yield (Fv/Fm) reached the maximal value (0.8) by T14, independent of light intensity (Figure 1D and Figure 1- figure supplement 1). This shows that PSII assembly, and more globally assembly of the photosynthetic machinery, occurs simultaneously with thylakoid membrane formation and that photosynthesis is operational almost immediately upon greening.

Our proteomic and lipidomic analyses suggest that chloroplast ultrastructural changes rely on specifically timed molecular changes. Proteomic analysis revealed the accumulation patterns of more than 5,000 unique proteins at eight time points during de-etiolation. These data provide information for plastid development and more widely on light-regulated developmental processes (Supplemental Dataset 1). Our dataset is more exhaustive regarding temporal resolution and the number of unique proteins detected than that of previous reports on chloroplast differentiation and de-etiolation (Bräutigam and Weber, 2009; Plöscher et al., 2011; Reiland et al., 2011; Wang et al., 2006).

Here, we focused on chloroplast-localized proteins, specifically on thylakoid membrane proteins. According to the SUBA4 localization consensus, 1,112 proteins were assigned to plastids, which covers about a third of the total plastid proteome (Ferro et al., 2003; Hooper et al., 2017; Kleffmann et al., 2007). Our data suggest that the reorganisation of pre-existing molecules rather than *de novo* synthesis is responsible for the major chloroplast ultrastructural changes that occur between T0 and T4. These results are consistent with other studies reporting only minor increases in protein accumulation and translation during initial chloroplast differentiation (Dubreuil et al., 2018; Kleffmann et al., 2007; Reiland et al., 2011). GO analysis combined with expression pattern–based hierarchical clustering highlighted that most photosynthesis-related proteins are globally coregulated (Figure 5- figure supplement 1, clusters 2 and 6). However, targeted immunoblot analysis revealed different accumulation dynamics for specific photosystem subunits: PSI subunits were detected at later time points than PSII subunits, but thereafter PSI subunit accumulation was faster (Figure 6). The kinetics of different photosynthetic parameters were consistent with the sequential activation of PSII and PSI, in particular photochemical quenching, which showed increased oxidation of the plastoquinone pool by T14 (Figure 1- figure supplement 1). Early accumulation of proteins such as Lhcb5, −6, and PSBS could be a way to quickly induce photoprotective mechanisms such as non-photochemical quenching to prevent PSII photodamage during initial photosynthetic machinery assembly. Differences in PSI and PSII accumulation dynamics and activity have been consistently observed in other chloroplast development experimental systems, including in Arabidopsis cell cultures, during germination and development of Arabidopsis seedlings in the light, and in tobacco leaves upon reillumination after dark adaptation (Armarego-Marriott et al., 2019; Dubreuil et al., 2018; Liang et al., 2018). The molecular mechanisms underlying this differential accumulation are currently unknown; however, preferential localization of the PSI and PSII protein complexes in specific thylakoid membrane domains (lamellae and grana, respectively) and the time taken to establish these domains during chloroplast development (i.e. grana appear later than stromal lamellae) may play influential roles.

Chloroplast membranes have a specific composition that differs from that of other cell membranes. Galactolipids constitute the bulk of the thylakoid membranes, but are mostly absent from other membrane systems under growth conditions where phosphorus nutrient is available (Jouhet et al., 2007). MGDG and DGDG represent around 80% of the thylakoid membrane lipids. The absolute quantification of 12 types of MGDG and DGDG galactolipids (representing the major forms) revealed specific patterns of accumulation (Figure 7). Results showed a gradual accumulation of MGDG and DGDG galactolipids derived from the ER pathway from T8 to T24, whereas galactolipids from the PL pathway started to accumulate after one day of light exposure (T24). This illustrates the different galactolipid compositions of etioplasts and chloroplasts: ER-pathway galactolipids are predominant in the etioplast whereas PL-pathway galactolipids are predominant in the chloroplast. As no significant changes in lipid accumulation were observed by T4, it appears likely that the emergence of PTs relies on the existing lipids in the etioplast PLB, as suggested also by Armarego-Marriott et al. (2019). At later time points, galactolipids from both the ER and PL pathways constitute the lipid matrix of the thylakoid membrane. How the two galactolipid biosynthesis pathways are regulated during development and/or upon light treatment remains to be elucidated; however, we hypothesize that the PL pathway gains traction after T24 when photosynthetic capacity is fully established.

### Chloroplast development: ‘Chloroplast Proliferation Phase’

Chloroplast development continued between T24 and T96, during which thylakoid membranes acquired grana stacks with more clearly defined organisation (Figure 2). Thylakoid surface increased by only 41%; however, chloroplasts continued to enlarge at a rate comparable to previous de-etiolation stages (T0–T24). This chloroplast volume expansion may be caused by enlargement of extra-thylakoidal spaces occupied by emerging starch granules. These results suggest that large amounts of lipids and proteins are necessary to build up the thylakoid membrane until T24, whereas increases in lipids and proteins between T24 and T96 enable the expansion of already functional thylakoid membranes in preparation for chloroplast division. Indeed, chloroplast number per cell increased during de-etiolation, a process that depends on the division of pre-existing chloroplasts.

Both chloroplasts and mitochondria divide through the activity of supramolecular complexes that constitute the organelle division machineries (Yoshida, 2018). As chloroplast proliferation was observed between T24 and T96, chloroplast division may correlate with developmental stage of the organelle. Components of the chloroplast division machinery (e.g. FtsZ and ARC5) were detectable in etioplasts; however, their protein levels accumulated significantly during de-etiolation as chloroplasts proliferated (Figure 8C and D). Interestingly, the capacity to divide appeared to correlate with a minimum chloroplast volume of about 100 μm^3^, even at T24 when most chloroplasts were smaller (Figure 8E and Figure 4B). Whether and how chloroplast size and developmental stage can be sensed to activate the chloroplast division machinery remains poorly understood and requires further study.

### A model of thylakoid expansion

Our mathematical model describing the expansion of thylakoid surface per seedling over time considered the surface area occupied by the membrane lipids MGDG and DGDG and the major photosynthetic complexes PSII, PSI, and Cyt *b*_6_*f*. We omitted some components that contribute to the total thylakoid membrane surface (e.g. the protein complexes ATP synthase and NDH, and the lipid sulfoquinovosyldiacylglycerol; together grouped as ‘ε’ in Equation 2). The predictions made by our model fit the surface estimated by SBF-SEM at T4 and T24, whereas they do not fit that at T96. This means that compounds used to generate the mathematical model appear to contribute most to changes in thylakoid surface during early stages of de-etiolation (the structure establishment phase). By contrast, during the later stages of de-etiolation (the chloroplast proliferation phase), the contribution of other compounds omitted in our model is obviously required to build up thylakoid surface.

Our proteomics data (Figure 5- figure supplement 1 and Dataset 2) revealed some proteins that increased between T24 and T96, such as the FtsH protease (AT2G30950). FtsH proteases have a critical function during thylakoid biogenesis. In Arabidopsis, they constitute a hetero-hexameric complex of four FtsH subunits, which is integrated in the thylakoid membrane (Kato and Sakamoto, 2018). Although the FtsH complex surface area is unknown in Arabidopsis, it can be considered as a potential compound contributing to the thylakoid surface changes missing from our mathematical model. Other proteins, such as those involved in carotenoid biosynthesis (AT3G10230) or fatty acid metabolism (AT1G08640), also increased significantly after T24, implying that they contribute to the ‘ε’ factor.

A follow-up study would be to test the model under different conditions to investigate how this biological system responds to internal (perturbing hormone concentrations, genetic modification of thylakoid lipid and protein composition) or external (different qualities of light) factors. This could be instrumental in revealing new potential regulatory mechanisms of thylakoid biogenesis and maintenance.

Upon de-etiolation, the development of photosynthetic capacity relies on successful chloroplast biogenesis. At the cellular level, this process is expected to be highly coordinated with the metabolism and development of other organelles. Lipid synthesis involves lipid exchanges between chloroplasts and the endoplasmic reticulum. How lipid trafficking is organised remains poorly understood, but could require membrane contact sites between these two organelles (Michaud and Jouhet, 2019). Physical interaction between mitochondria and chloroplasts have been reported previously in diatoms (Bailleul et al., 2015; Flori et al., 2017). Whether such contact sites occur and are functional in plants is unknown; however, these mechanisms are hypothesized to exist since it is necessary that chloroplasts exchange metabolites with mitochondria and peroxisomes to ensure activation of photorespiration concomitantly with photosynthesis. The study of membrane contact sites is an emerging field in cell biology (Scorrano et al., 2019). Future work will focus on analysing the dynamics and functionality of contact sites between chloroplast membranes and other organelles, and investigate the general coordination of plant cell metabolism during de-etiolation. These questions could be further addressed using the SBF-SEM stacks and proteomic resource described here.

## Materials and methods

### Plant material and Growth conditions

*Arabidopsis thaliana* seeds (Columbia ecotype) were surface-sterilized with 70% (v/v) ethanol with 0.05% (v/v) Triton X-100, then washed with 100% ethanol. Seeds were sown on agar plates containing 0.5 × Murashige and Skoog salt mixture (Duchefa Biochemie, Haarlem, Netherlands) without sucrose. Following stratification in the dark for 3 days at 4°C, seeds were irradiated with 40 μmol m^−2^ s^−1^ for 2 h at 21°C and then transferred to the dark for 3 days growth at 21°C. Etiolated seedlings were collected in the dark (0 h of light; T0) and at selected time points (T4, T8, T12, T24, T48, T72, T96) upon continuous white light exposure (40 μmol m^−2^ s^−1^ at 21°C).

### Photosynthetic parameters

Maximum quantum yield of photosystem II (ΦMAX= F_V_/F_M_ =(Fm−Fo)/Fm where Fm is the maximal fluorescence in dark adapted state, Fo is minimal fluorescence in dark adapted state, Fv is the variable fluorescence (Fm−Fo)), photosystem II quantum yield in the light (ΦPSII), and photochemical quenching (qP) were determined using a Fluorcam (Photon Systems Instruments) with blue-light LEDs (470 nm). Plants were dark adapted for a minimum of 5 min before measurement.

### Chlorophyll concentration

Chlorophylls were extracted in 4 volumes of dimethylformamide (DMF) (v/w) overnight at 4°C. After centrifugation, chlorophylls were measured using a NanoDrop™ instrument at 647 nm and 664 nm. Chlorophyll contents were calculated according to previously described methods (Porra et al., 1989).

### Transmission electron microscopy (TEM)

Samples were fixed under vacuum (200 mBar) in 0.1 M cacodylate buffer (pH 7.4) containing 2.5% (w/v) glutaraldehyde and 2% (w/v) formaldehyde (fresh from paraformaldehyde) for 4 h and left in the fixation solution for 16 h at 4°C. Samples were then incubated in a solution containing 3% (w/v) potassium ferrocyanide and 4 mM calcium chloride in 0.1 M cacodylate buffer combined with an equal volume of 4% (w/v) aqueous osmium tetroxide (OsO_4_) for 1 h, on ice. After the first heavy metal incubation, samples were rinsed with ddH_2_O and treated with 1% (w/v) thiocarbohydrazide solution for 1 h at 60°C. Samples were rinsed (ddH_2_O for 15 min) before the second exposure to 2% (w/v) OsO_4_ aqueous solution for 30 min at room temperature. Following this second exposure to osmium, tissues were placed in 1% (w/v) uranyl acetate (aqueous) and left overnight at 4°C. The samples were rinsed with ddH_2_O for 15 min, and placed in the lead aspartate solution for 30 min at 60°C. Samples were dehydrated in a series of aqueous ethanol solutions ranging from 50% (v/v) to 100%, then embedded in Durcupan resin by successive changes of Durcupan resin/acetone mixes, with the last imbibition in 100% Durcupan resin. Polymerization of the resin was conducted for 48 h at 60°C (Deerinck et al., 2010). Ultra-thin sections (70 nm) were cut using Ultrathin-E microtome (Reichert-Jung) equipped with a diamond knife. The sections were analysed with a Philips CM-100 electron microscope operating at 60 kV.

### Confocal microscopy

To derive the chloroplast and cell volumes, images of 1–5-μm thick sections of cotyledon cells were acquired with X10 and X40 oil immersion objectives using a LEICA TCS SP5 confocal laser scanning microscope. Chlorophyll was excited using a red laser (33%) and spectral detection channel was PMT3.

### SBF-SEM

SBF-SEM was performed on Durcupan resin–embedded cotyledons representing the four de-etiolation time points T0, T4, T24, and T96. Overview of the mesophyll tissue (≈600 images) and zoomed stacks of the chloroplasts (≈300 images) were acquired. Voxel size of T4 zoomed stacks: 3.9 × 3.9 × 50 nm; T24: 4.7 × 4.7 × 50 nm; T96: 5.6 × 5.6 × 50 nm. Voxel size for T0 overview: 9.5 × 9.5 × 100 nm; T4: 19.3 × 19.3 × 100 nm; T24: 40 × 40 × 200 nm; T96: 43.5 × 43.5 × 200 nm.

Acquired datasets were aligned and smoothed respectively, using the plugins MultiStackReg and 3D median filter, provided by the open-source software Fiji.

We performed a stack-reslice from Fiji to generate a new stack by reconstructing the slices at a new pixel depth to obtain isotropic voxel size and improve z-resolution. The segmentation and 3D mesh geometry information of plastid /thylakoid (T0, T4, T24 and T96) were implemented by open-source software 3D Slicer (Fedorov et al., 2012) and MeshLab (Cignoni et al., 2008) respectively.

### Segmentation, 3D reconstruction, and surface and volume quantification

Segmentation and 3D reconstruction of 3View and confocal images were performed using Amira software (FEI Visualization Sciences Group). Specifically, prolamellar body, thylakoids, and envelope membranes as well as the cells were selected using a semi-automatic tool called Segmentation Editor. From the segmented images, triangulated 3D surfaces were created using Generate Surface package. Quantification of morphometric data (Area 3D and volume 3D) was acquired using Label Analysis package.

### Analysis of grana segmentation

Grana structures acquired from SBF-SEM were selected in Amira. The grana selections were converted in line set view in Amira software using the Generate Contour line package. To complete the grana segmentation, the line set views were imported into the Rhino 6 software (Robert McNeel & Associates, USA). Every granum was segmented in layers with a specific thickness and distance according to quantitative data collected (Figure 2- figure supplement 1 and Figure 3- figure supplement 1). After segmentation, images were re-imported in Amira software to quantify perimeter using the Label Analysis package.

### Chloroplast number determination

Chloroplasts per cell were counted manually using Image J software (Wayne Rasband, National Institutes of Health). From the same SBF-SEM stack, 5 and/or 6 cells were cropped at each time point (T0, T4, T24, and T96) to quantify chloroplast number per cell. From TEM images, chloroplast number/cell was determined at T24 (16 cells), T48 (12 cells), T72 (12 cells), and T96 (17 cells). TEM images were acquired from two independent experiments.

### Liquid chromatography–mass spectrometry analysis and protein quantification

Etiolated seedlings were grown as described above. At each time point, ca. 80 seedlings were pooled, frozen in liquid nitrogen, and stored at −80°C until use. Frozen material was ground with a mortar and pestle, and 40–80 mg of plant material was used for protein and peptide preparation using the iST kit for plant tissues (PreOmics, Germany). Briefly, each sample was resuspended in 100 μL of the provided ‘Lysis’ buffer and processed with High Intensity Focused Ultrasound (HIFU) for 1 min by setting the ultrasonic amplitude to 65% to enhance solubilization. For each sample, 100 μg of protein was transferred to the cartridge and digested by adding 50 μL of the provided ‘Digest’ solution. After 180 min of incubation at 37°C, the digestion was stopped with 100 μL of the provided ‘Stop’ solution. The solutions in the cartridge were removed by centrifugation at 3,800 *g*, whereas the peptides were retained on the iST filter. Finally, the peptides were washed, eluted, dried, and re-solubilized in 18.7 μL of solvent (3% (v/v) acetonitrile, 0.1% (v/v) formic acid).

Mass spectrometry (MS) analysis was performed on a Q Exactive HF-X mass spectrometer (Thermo Scientific) equipped with a Digital PicoView source (New Objective) and coupled to a M-Class UPLC (Waters). Solvent composition at the two channels was 0.1% (v/v) formic acid for channel A and 0.1% formic acid, 99.9% (v/v) acetonitrile for channel B. For each sample, 2 μL of peptides were loaded on a commercial MZ Symmetry C18 Trap Column (100 Å, 5 μm, 180 μm × 20 mm, Waters) followed by nanoEase MZ C18 HSS T3 Column (100 Å, 1.8 μm, 75 μm × 250 mm, Waters). The peptides were eluted at a flow rate of 300 nL/min by a gradient of 8–27% B in 85 min, 35% B in 5 min, and 80% B in 1 min. Samples were acquired in a randomized order. The mass spectrometer was operated in data-dependent mode (DDA), acquiring a full-scan MS spectra (350−1400 m/z) at a resolution of 120,000 at 200 m/z after accumulation to a target value of 3,000,000, followed by HCD (higher-energy collision dissociation) fragmentation on the 20 most intense signals per cycle. HCD spectra were acquired at a resolution of 15,000 using a normalized collision energy of 25 and a maximum injection time of 22 ms. The automatic gain control (AGC) was set to 100,000 ions. Charge state screening was enabled. Singly, unassigned, and charge states higher than seven were rejected. Only precursors with intensity above 250,000 were selected for MS/MS. Precursor masses previously selected for MS/MS measurement were excluded from further selection for 30 s, and the exclusion window was set at 10 ppm. The samples were acquired using internal lock mass calibration on m/z 371.1012 and 445.1200. The mass spectrometry proteomics data were handled using the local laboratory information management system (LIMS) (Türker et al., 2010).

Protein quantification based on precursor signal intensity was performed using ProgenesisQI for Proteomics (v4.0.6403.35451; nonlinear dynamics, Waters). Raw MS files were loaded into ProgenesisQI and converted to mzln files. To select the alignment reference, a group of samples that had been measured in the middle of the run (to account for drifts in retention times) and derived from de-etiolation time point T12 or later (to account for increasing sample complexity) was preselected, from which replicate 3 of time point T48 was then automatically chosen as best alignment reference. After automatic peak picking, precursor ions with charges other than 2+, 3+, or 4+ were discarded. The five highest-ranked MS/MS spectra, at most, for each peptide ion were exported, using the deisotoping and charge deconvolution option and limiting the fragment ion count to 200 peaks per MS/MS. The resulting Mascot generic file (.mgf) was searched with Mascot Server version 2.6.2 (www.matrixsicence.com) using the following settings: trypsin digest with up to two missed cleavages allowed; carbamidomethylation of cysteine as fixed modification; N-terminal acetylation and oxidation of methionine residue as variable modifications; precursor ion mass tolerance 10 ppm; fragment ion (MS/MS) tolerance 0.04 kDa. This search was performed against a forward and reverse (decoy) Araport11 database that included common MS contaminants and iRT peptides. The mascot result was imported into Scaffold Q+S (v4.8.9; Proteome Software Inc), where a spectrum report was created using a false discovery rate (FDR) of 10% and 0.5% at the protein and peptide level, respectively, and a minimum of one identified peptide per protein. After loading the spectrum report into ProgenesisQI, samples were normalized using the “normalize to all proteins” default settings (i.e. normalization was performed to all ions with charges 2+, 3+ or 4+). Samples were grouped according to de-etiolation time point in a between-group analysis with 4 replicates for each condition, except for time point T0 and T48, where n = 3. For these two time points, one replicate each had been discarded it appeared as an outlier in principal component analysis (PCA) of protein abundances between different runs (Supplemental dataset 1).. Quantification employed the Hi-N method, measuring the three most abundant peptides for each protein (Grossmann et al., 2010), and associated statistics (q-value, PCA etc.) were calculated in ProgenesisQI. Quantification also used protein grouping, which assigns proteins for which only shared but no unique peptides were identified to a ‘lead’ identifier containing all these shared peptides and thus having the greatest coverage among all grouped identifiers or highest score where coverage is equal. Quantification was restricted to protein (groups) with at least two identified peptides among which at least one is unique to the protein (group). Using these requirements, 5082 Arabidopsis proteins (or groups) were identified. Since 13 additional identifications were exclusively associated with decoy proteins, the false discovery rate at the protein level is estimated to be 0.3%. .

### Immunoblot analysis

Proteins were extracted from whole seedlings in 4 volumes (w/v) of SDS-PAGE sample buffer (0.2 M Tris/HCL pH 6.8, 0.4 M dithiothreitol, 8% (w/v) SDS, 0.4% (w/v) Bromophenol blue, and 40% (v/v) glycerol).

Proteins were denatured for 15 min at 65°C and cell debris were removed by centrifugation for 5 min at 16,000 *g*. Proteins were separated on SDS-PAGE (10–15% (w/v) polyacrylamide concentrations depending on the molecular weight of the protein of interest) and transferred onto a nitrocellulose membrane for immunoblotting (overnight at 4°C) in Dunn buffer (10 mM NaHCO_3_, 3 mM Na_2_CO_3_, 0.01% (w/v) SDS, and 20% ethanol).

Absolute quantification of PsbA, PetC, and PsaC was performed according to Agrisera instructions and using recombinant proteins (PsbA AS01 0116S, PetC AS08 330S, and PsaC AS04 042S; Agrisera, Vännäs, SWEDEN). Three respective calibration curves for the three recombinant proteins were created. Concentrations used to generate the PsbA and PetC calibration curves were 1.75, 2.5, 5, and 10 (ng/μL). Concentrations used to generate the PsaC calibration curve were 0.375, 0.75, 1.5, and 3 (ng/μL). Immunodetections were performed using specific antibodies: anti-Actin (Sigma, A0 480) at 1/3,000 dilution in 5% (w/v) milk in Tris-buffered saline (TBS); anti-Lhcb2 (Agrisera, AS01 003), anti-D1(PsbA) (Agrisera, AS05 084), anti-PsbO (Agrisera, AS14 2825), anti-PsbD (Agrisera, AS06 146), anti-PetC (Agrisera, AS08 330), and anti-AtpC (Agrisera, AS08 312) at 1/5,000 dilution in 5% milk/TBS; Anti-PsaD (Agrisera, AS09 461) at 1/2,000 in 5% milk/TBS; and anti-PsaC (Agrisera, AS042P) and anti-ARC5 (Agrisera, AS13 2676) at 1/2,000 in 3% (w/v) bovine serum albumin (BSA) in TBS. Anti-FtsZ-1 and anti-FtsZ2-1/FtsZ 2-2 (El-Shami et al., 2002; Karamoko et al., 2011) and were used at 1/2,000 dilution in 5% milk/TBS. After incubation with primary antibodies overnight at 4°C, blots were washed 3 times in TBS containing 0.1% (v/v) Tween without antibodies for 10 minutes and incubated for 1 h at RT with horseradish peroxidase–conjugated secondary antibodies (1/3,000 (v/v) anti-rabbit or anti-mouse secondary antibodies, Agrisera). Chemiluminescence signals were generated with Enhanced chemiluminescence reagent (1 M Tris/HCl pH 8.5, 90 mM coumaric acid, and 250 mM luminol) and detected with a Fujifilm Image – Quant LAS 4000 mini CCD (GE Healthcare). Quantifications were performed with ImageQuant TL software (GE Healthcare).

### Lipid profiling

Lipids were extracted from whole seedlings ground in a mortar and pestle under liquid nitrogen. Ground plant material corresponding to 40–80 mg fresh weight was suspended in tetrahydrofuran:methanol (THF/MeOH) 50:50 (v/v). 10–15 glass beads (1 mm in diameter) were added followed by homogenization (3 min, 30 Hz,) and centrifugation (3 min, 14 000 *g*, at 4°C). The supernatant was removed and transferred to an HPLC vial. Lipid profiling was carried out by ultra-high pressure liquid chromatography coupled with atmospheric pressure chemical ionization-quadrupole time-of-flight mass spectrometry (UHPLC-APCI-QTOF-MS) (Martinis et al., 2011). Reverse-phase separation was performed at 60°C on an Acquity BEH C18 column (50 × 2.1 mm, 1.7 μm). The conditions were the following: solvent A = water; solvent B = methanol; 80–100% B in 3 min, 100% B for 2 min, re-equilibration at 80% B for 0.5 min. Flow rate was 0.8 ml min^−1^ and the injection volume 2.5 μl. Data were acquired using MassLynx version 4.1 (Waters), and processed with MarkerLynx XS (Waters). Peak lists consisting of variables described by mass-to-charge ratio and retention time were generated (Martinis et al., 2011; Spicher et al., 2016).

Absolute quantification of mono-(MGDG) and di-galactosyldiacylglycerol (DGDG) was conducted by creating calibration curves using MGDG (reference number 840523) and DGDG (reference number 840523) products of Avanti Company. Calibration curves were prepared using the following concentrations: 0.08, 0.4, 2, 10, and 50 μg ml^−1^ of MGDG or DGDG.

### Mathematical Model

A non-linear mixed effects model (with fixed effect of time and random effect of replicates on 3 of the parameters), built on a 4-parameter logistic function, was implemented in R (free software created by Ross Ihaka and Robert Gentleman, Auckland University, New Zealand), following the examples in Pinheiro and Bates (2000). The R-packages used are: nlme (Pinheiro and Bates, 2000), effects, lattice and car (Fox and Weisberg, 2018). To account for self-correlation at the replicate level, we proceeded to fit an overall mixed-effects model to the data (package ‘nlme’ from R), using the replicate’s as random effect term (Figure 9_supplement 1). The four parameters *a, b, c* and *d* have been calculated (Figure 9_supplement 1) and the three plots (one for each biological replicate) (Figure 9_supplement 1) indicated the fitting curve for a series of data points.

## Supporting information

Supplemental Datset 1

## Acknowledgements

This work was supported by the University of Neuchâtel and ETH Zurich, a grant from the Swiss National Science Foundation (3100A0-112638) to E.D., and grants 31003A_156998 and 31003A_176191 to F.K.. We thank Jonas Grossmann, Laura Kunz and Paolo Nanni from the Functional Genomic Center Zurich (FGCZ) for peptide preparation for mass spectrometry, acquisition of the raw data and help with associated data analysis, the ETH Zurich microscopy facility (ScopeM) for advice in conducting SBF-SEM analysis. We thank Slobodeanu Radu Alexandru and Federico Giacomarra for help with bioinformatics analysis, and Romain Bessire for help with image processing software. We thank Roman Ulm and Michel Goldschmidt-Clermont for critical reading of the manuscript.

## Author contributions

Conceptualization, R.P., T.P., S.Z., F.K., and E.D.; Investigation, R.P., S.E., B.P., D.F., C.U., G.G., and E.D.; Writing R.P, B.P., F.K. and E.D.; Supervision, F.K. and E.D.

## Competing interests

The authors declare no competing interests.

## Figure Supplements

**Figure 1- figure supplement 1: Photosynthesis parameters during de-etiolation**. Maximum photosynthetic quantum yield of PSII (Fv/Fm) of plants (dark-adapted for 5 minutes) grown under different light intensities (A). Photochemical quenching (B) and efficiency of the photosystem PSII (Φ PSII; C) measurements were made on 3-day-old etiolated seedlings that were de-etiolated under continuous light (40 μmol/m^2^/s) using a Fluorcam (Photon System Instrument). Error bars indicate ± SD (n=10).

**Figure 2- figure supplement 1: Measurement of lamella thickness**. (A) TEM chloroplast micrographs of 3-day-old, dark-grown *Arabidopsis thaliana* (Columbia) seedlings illuminated for 96 h in continuous white light (40 μmol/m^2^/s) were used to measure the thickness of lamellae that constitute the grana stack. Measurements were performed using ImageJ. Scale bar: 100 nm. (B) Equation used to calculate the thickness of one lamella. (C) Data indicate mean ± SD (n=10 for 2 lamellae and n=7 for 3 lamellae).

**Figure 4- figure supplement 1: Grana segmentation (T24)**. (A) Selection of thylakoid membrane exposed to the stroma was acquired using Amira. (B) The perimeter of the grana structures showed in black were segmented in layers of a specific thickness and distance using Rhino software, with the corresponding thickness (lamellae and stromal gap) measured and calculated as described in Figure 2- figure supplement 1. Grana segmentation was performed using thylakoid membrane of de-etiolating seedlings exposed to continuous white light (40 μmol/m^2^/s) for 24 (T24) and 96 (T96) h. A representative example of a T24 replicate is illustrated here. (C) Schematic representation of the grana stack perimeter comprising margins, end membranes, and intergranal lamellae. (D) Equation used to calculate the percentage of the grana stack surface area relative to total thylakoid surface area.

**Figure 5- figure supplement 1: Accumulation dynamics of selected plastid proteins during de-etiolation**. Hierarchical clustering (Euclidean, average linkage) of normalized protein abundance (log2 fold changes) for plastid-localized proteins during de-etiolation. Normalization was performed to the last time point (96 h). Defined clusters are indicated with different colours (1= purple; 2= pink; 3=turquoise; 4= brown; 5 = light green; 6 = dark green). Protein IDs (AGI) and names are legible upon zoom-in.

**Figure 6- figure supplement 1: Quantification of photosynthesis-related proteins**. (A) Immunodetection of PsbA, PetC, and PsaC during de-etiolation. Dilutions were used for the later time points to avoid saturation of the signal. (B) Different bands were detected by Amersham Imager program and quantified by Image QuantTL (Amersham). (C) Calibration curves were created using recombinant proteins (Agrisera). Calibration curve composition: PsbA 10 ng (A; lane a), 5 ng (b), 2.5 ng (c), and 1.25 ng (d); PetC 10 ng (e), 5 ng (f), 2.5 ng (g), and 1.25 ng (h); PsaC 3 ng (i), 1.5 ng (l), 0.75 ng (m), and 0.325 ng (n). The analysis was carried out on 3–4 independent experiments (BIO1–4).

**Figure 8- figure supplement 1. Chloroplast proliferation in parallel with cell expansion**. SEM micrographs of 3-day-old, dark-grown *Arabidopsis thaliana* (Columbia) seedlings illuminated for 0 h (T0; A), 4 h (T4; B), 24 h (T24; C), and 96 h (T96; D) in continuous white light (40 μmol/m^2^/s). Palisade (PA) and spongy (SP) cells are indicated. Scale bars: 15 μm. (E) 3D reconstruction of a palisade cell at T24 after segmentation of chloroplasts and cell plasma membrane. (F–I) Confocal images of cotyledons of dark-grown seedlings at T24 (F), T48 (G), T72 (H), and T96 (I). Scale bars: 10 μm. (L–O) TEM micrographs of cotyledon cells of dark-grown seedlings at T24 (L), T48 (M), T72 (N), and T96 (O). L–M, scale bars: 2 μm; N– O, scale bars: 5 μm. (P) Cell perimeter measured with Amira software using (red line). The Z-depth of each stack corresponds to 1 μm. Relative chloroplast number per cell was counted using 2D TEM images (black line). Red error bars indicate ± SD (n= 17). Black error bars indicate ± SD (n= 3–4). (Q) Box plots of single chloroplast total volume quantified at T48 and T72. Each box corresponds to the distribution of a chloroplast population analysed using confocal and SBF-SEM stacks. n=66 (T48); 62 (T72).

**Figure 9- figure supplement 1: Non-linear mixed effect model of thylakoid surface during de-etiolation**. (A) Total surface of thylakoid membrane components (in μm^2^) in function of de-etiolation time point. (B) Individual plots for each biological replicate. (C) Values, standard errors, t-value, and P-value of the four parameters (a, b, c, and d) used in the main equation. Smodel = surface of thylakoid at a specific time (t)

t = time of light exposure (h)

a = asymptote (to the left if c>0)

b= right asymptote (to the right if c>0)

c= proportional to the slope of the curve at the inflection point

d= inflection point (point at which the mean Smodel value is reached)

**Figure 9- figure supplement 2: Morphometric analysis of cotyledons**. (A) Cotyledon surface area of 3-day-old, dark-grown *Arabidopsis thaliana* (Columbia) seedlings illuminated with 24 h (T24) and 96 h (T96) of continuous white light (40 μmol/m^2^/s). (B) The thickness (T) of mesophyll tissue constituted of palisade (PA), spongy (SP) cells, and vascular system (VS) in addition to the epidermal tissue was measured. Error bars indicate ± SD (n=4). (C) Estimation of cotyledon volume. Error bars indicate ± SD (n=3). (D) Estimation of the number of cells per cotyledon (see Supplemental Dataset 4 for calculations)

## Notes

### Competing Interest Statement

The authors have declared no competing interest.

