## Supplementary material for "Two distinct phases of chloroplast biogenesis during de-etiolation in *Arabidopsis thaliana*": tables

| | Method | T0 | T4 | T8 | T12 | T24 | T48 | T72 | T96 |
|---|---|---|---|---|---|---|---|---|---|
| *Chloroplast volume (µm$^3$)* | SBF-SEM | **12.27** (± 2.3) | **9.4** (± 4.8) | — | — | **62** (± 2.04) | — | — | **112.14** (± 4.3) |
| *Thylakoid surface (µm$^2$)* | SBF-SEM | — | **67.0** (± 29.5) | — | — | **1476** (± 146) | — | — | **2086** (± 393) |
| *Grana lamellae / total thylakoid surface* | | — | — | — | — | **2.55** (± 0.11) | — | — | **2.08** (± 0.57) |
| *Thylakoid/envelope surface* | | — | **1,02** (± 0.15) | — | — | **7.37** (± 0.51) | — | — | **6.83** (± 1.40) |
| *Length of plastid (µm)* | TEM | **2** (± 0.90) | **2.8** (± 0.90) | — | — | **5.1** (± 1.47) | — | — | **6** (± 1.62) |
| *Stroma lamellae volume (µm$^3$)* | SBF-SEM | — | **2.43** (± 0.95) | — | — | **17.87** (± 1.04) | — | — | **29.17** (± 19.4) |
| *Chloroplast volume (µm$^3$)* | Confocal | — | — | — | — | **61.5** (± 11.2) | **70.1** (± 10.2) | **85** (± 22) | — |
| *Cell Volume (µm$^3$)* | 3 View | **1173** (± 284) | **1891** (± 362) | — | — | **6103** (± 1309) | — | — | **52597** (± 12671) |
| *Cell perimeter (µm)* | TEM | — | — | — | — | **55.26** (± 6.1) | **46.43** (± 5.3) | **71.68** (± 7.0) | **92.77** (± 11.1) |
| *Number of chloroplast per cell* | SBF-SEM | **22** (± 6) | **25** (± 8) | — | — | **26** (± 6) | — | — | **112** (± 29) |
| *Number of cell per seedling* | | — | — | — | — | **≈ 3000** | — | — | **≈ 3000** |
| *Protein/GLs surface* | | **0.19** (± 0.05) | **0.23** (± 0.04) | **0.34** (± 0.03) | **0.52** (± 0.07) | **0.80** (± 0.14) | **0.80** (± 0.17) | **0.78** (± 0.07) | **0.87** (± 0.25) |
| *GLs (nmol/seedling)* | Lipidomics | **0.31** (± 0.03) | **0.31** (± 0.02) | **0.32** (± 0.02) | **0.54** (± 0.02) | **0.67** (± 0.04) | **1.28** (± 0.12) | **1.84** (± 0.01) | **2.20** (± 0.09) |
| *PsbA (nmol/seedling)* | Immunodetection | **6.9$^{E}$-06** (± 1.8E-06) | **9.2$^{E}$-06** (± 1.7E-06) | **1.5$^{E}$-05** (± 7.1E-07) | **3.2$^{E}$-05** (± 4.3E-06) | **9.3$^{E}$-05** (± 2.1E-05) | **2.0$^{E}$-04** (± 6.2E-05) | **3.9$^{E}$-04** (± 4E-05) | **6.2$^{E}$-04** (± 1.7E-04) |
| *PsaC (nmol/seedling)* | Immunodetection | n.d | n.d | n.d | **1.6$^{E}$-05** (± 2.5E-06) | **7.3$^{E}$-05** (± 2.4-E05) | **1.1$^{E}$-04** (± 7.2-E05) | **1.7$^{E}$-04** (± 4.2-E05) | **2.3$^{E}$-04** (± 1E-04) |
| *PetC (nmol/seedling)* | Immunodetection | **2.7$^{E}$-05** (±7.8E-06) | **2.8$^{E}$-05** (± 9.8E-06) | **2.5$^{E}$-05** (± 4.5E-06) | **5.3$^{E}$-05** (± 2.2E-05) | **1.2$^{E}$-04** (± 4.1E-05) | **1.8$^{E}$-04** (± 3.4E-05) | **5.7$^{E}$-04** (± 1.8E-04) | **7.9$^{E}$-04** (± 3.7E-04) |

| | T0 | T4 | T8 | T12 | T24 | T48 | T72 | T96 |
|---|---|---|---|---|---|---|---|---|
| MGDG | **1.11E+07**<br>SD = +/- 3.64E+05 | **1.15E+07**<br>SD = +/- 1.01E+06 | **1.11E+07**<br>SD = +/- 1.10E+06 | **1.75E+07**<br>SD = +/- 1.79E+06 | **4.16E+07**<br>SD = +/- 4.27E+06 | **8.65E+07**<br>SD = +/- 5.96E+06 | **1.68E+08**<br>SD = +/- 8.86E+06 | **2.35E+08**<br>SD = +/- 1.94E+07 |
| DGDG | **3.64E+06**<br>SD = +/- 4.04E+05 | **4.23E+06**<br>SD = +/- 5.29E+05 | **4.10E+06**<br>SD = +/- 1.31E+05 | **6.26E+06**<br>SD = +/- 4.70E+05 | **1.32E+07**<br>SD = +/- 9.73E+05 | **2.32E+07**<br>SD = +/- 1.83E+06 | **3.97E+07**<br>SD = +/- 2.56E+06 | **5.48E+07**<br>SD = +/- 3.71E+06 |
| PSII | **2.04E+06**<br>SD = +/- 5.38E+05 | **2.74E+06**<br>SD = +/- 5.30E+05 | **4.40E+06**<br>SD = +/- 2.12E+05 | **9.91E+06**<br>SD = +/- 1.29E+06 | **2.75E+07**<br>SD = +/- 6.42E+06 | **6.06E+07**<br>SD = +/- 1.85E+07 | **1.15E+08**<br>SD = +/- 1.19E+07 | **1.83E+08**<br>SD = +/- 5.17E+07 |
| PSI | **0E+00**<br>SD = +/-0E+00 | **0E+00**<br>SD = +/- 0E+00 | **0E+00**<br>SD = +/- 0E+00 | **8.95E+05**<br>SD = +/- 4.49E+05 | **1.33E+07**<br>SD = +/- 4.31E+06 | **2.10E+07**<br>SD = +/- 1.30E+07 | **3.04E+07**<br>SD = +/- 7.55E+06 | **4.24E+07**<br>SD = +/- 1.89E+07 |
| Cyt $b_6 f$ | **7.99E+05**<br>SD = +/- 2.33E+05 | **8.43E+05**<br>SD = +/- 2.91E+05 | **7.50E+05**<br>SD = +/- 1.33E+05 | **1.57E+06**<br>SD = +/- 6.71E+05 | **3.44E+06**<br>SD = +/- 1.22E+06 | **5.30E+06**<br>SD = +/- 1.01E+06 | **1.69E+07**<br>SD = +/- 5.48E+06 | **2.37E+07**<br>SD = +/- 1.11E+07 |

| | *surface in nm$^2$* | *reference* |
|---|---|---|
| MGDG | 0.82 | *Bottier et al., 2007* |
| DGDG | 0.64 | *Bottier et al., 2007* |
| PSII -LHCII ($C_2S_2M_2$) | 494 | *Caffarri et al., 2014* |
| cyt $b_6f$ | 49.5 | *Kurisu et al., 2003* |
| PSI | 300 | *Caffarri et al., 2014* |
