## Supplemental figures for "Two distinct phases of chloroplast biogenesis during de-etiolation in *Arabidopsis thaliana*"

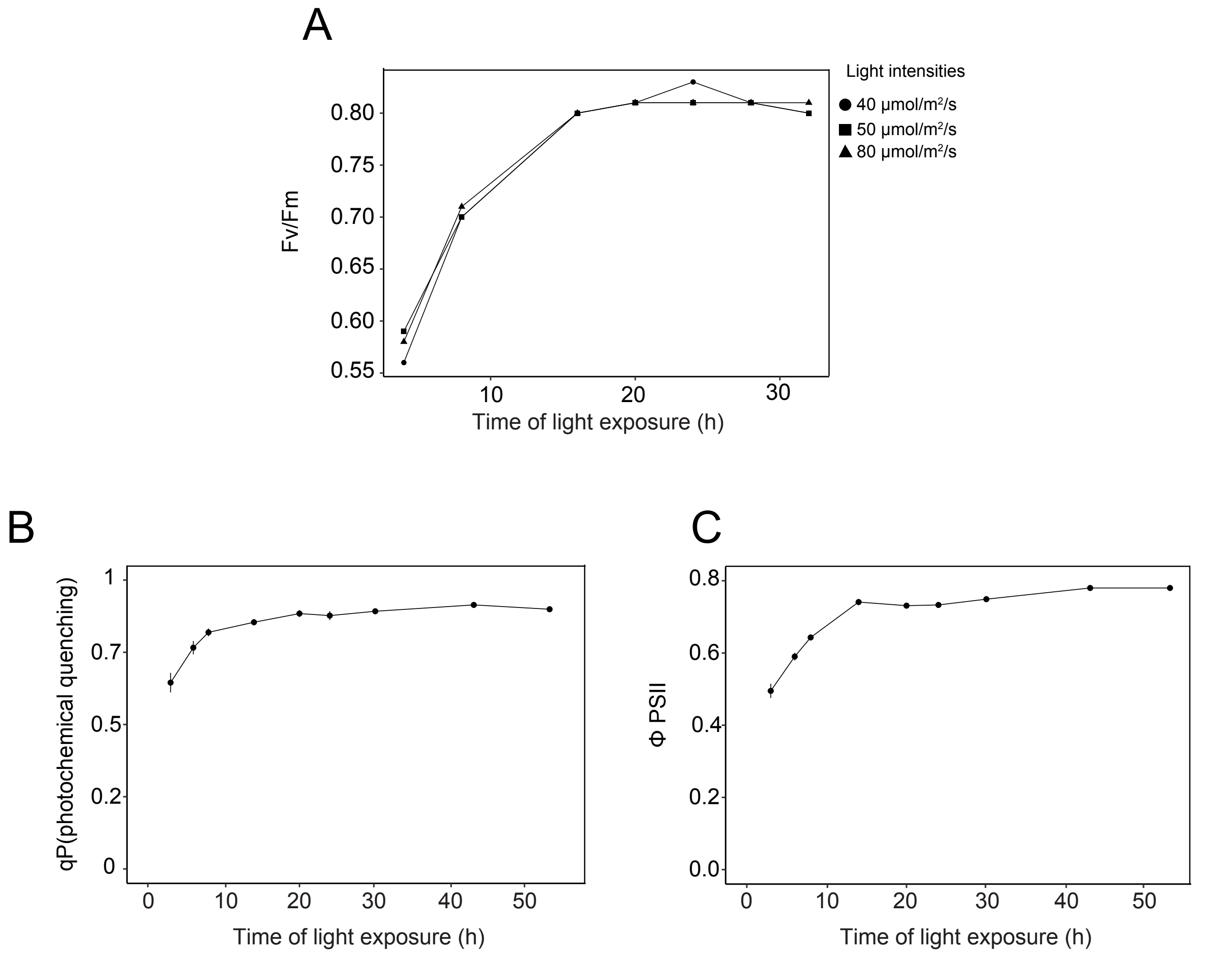
A
Light intensities
40 μmol/m²/s
50 μmol/m²/s
80 μmol/m²/s
Fv/Fm
0.55
0.60
0.65
0.70
0.75
0.80
10
20
30
Time of light exposure (h)
B
qP(photochemical quenching)
0
0.2
0.5
0.7
1
0
10
20
30
40
50
Time of light exposure (h)
C
Φ PSII
0.0
0.2
0.4
0.6
0.8
0
10
20
30
40
50
Time of light exposure (h)

A

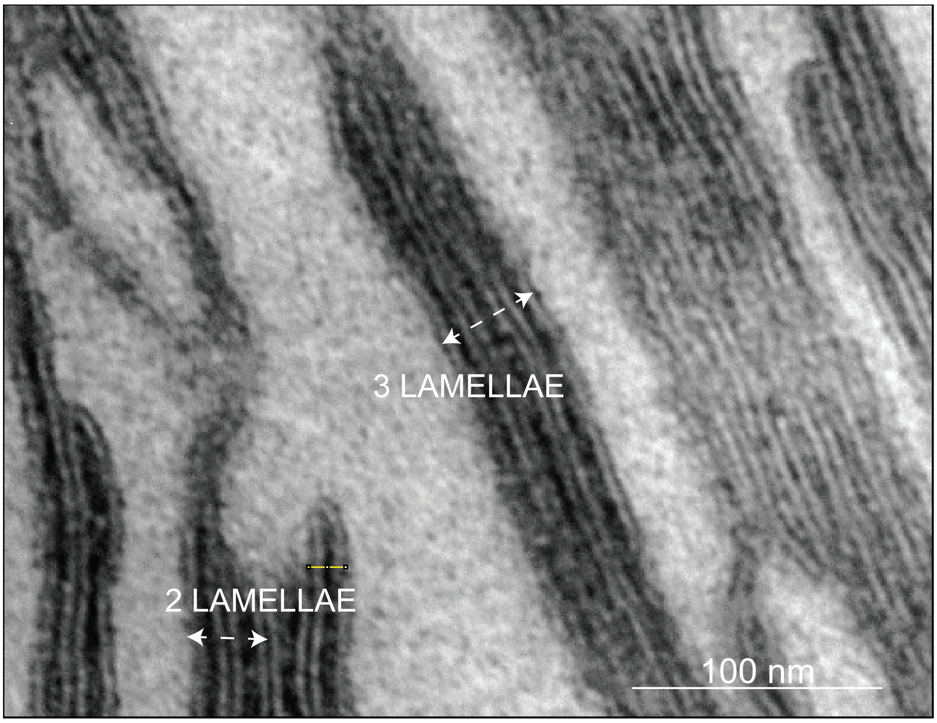


B

$$\frac{\textit{Thickness of 2 or 3 lamellae - (Stromal gap thickness * } \text{nb)}}{\textit{nb.lamellae measured}} = \textit{one lamellae thickness}$$

C

| | Thickness 2 lamellae (nm) | Thickness 3 lamellae (nm) | Stromal gap thickness *(Daum et al.,2010)* |
|---|---|---|---|
| T24 | 29.7 + / - 1.9 | 48.3 + / - 4.8 | |
| one lamellae thickness (T24) | $\frac{29.7-3.2}{2} = 13.2$ | $\frac{48.3-(3.2\quad 2)}{3} = 13.9$ | |
| T96 | 34.7 + / - 0.71 | 56 + / - 2.1 | 3.2 nm +/- 0.7 |
| one lamellae thickness (T96) | $\frac{34.7-3.2}{2} = 15.7$ | $\frac{56-(3.2\quad 2)}{3} = 16.5$ | |

.

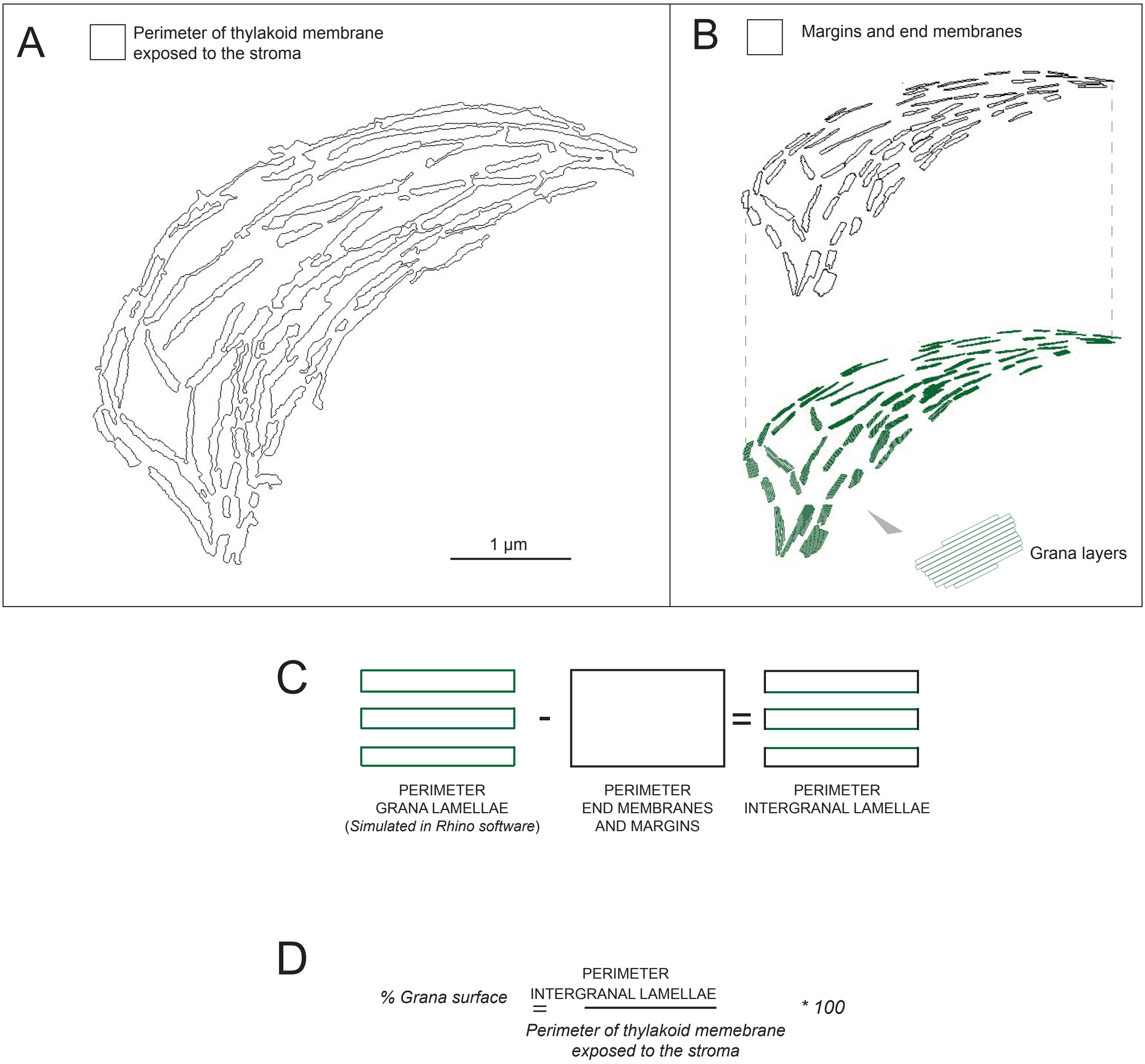

A
Perimeter of thylakoid membrane exposed to the stroma
1 µm
B
Margins and end membranes
Grana layers
C
PERIMETER GRANA LAMELLAE (Simulated in Rhino software)
-
PERIMETER END MEMBRANES AND MARGINS
=
PERIMETER INTERGRANAL LAMELLAE
D
% Grana surface = PERIMETER INTERGRANAL LAMELLAE / Perimeter of thylakoid memebrane exposed to the stroma * 100

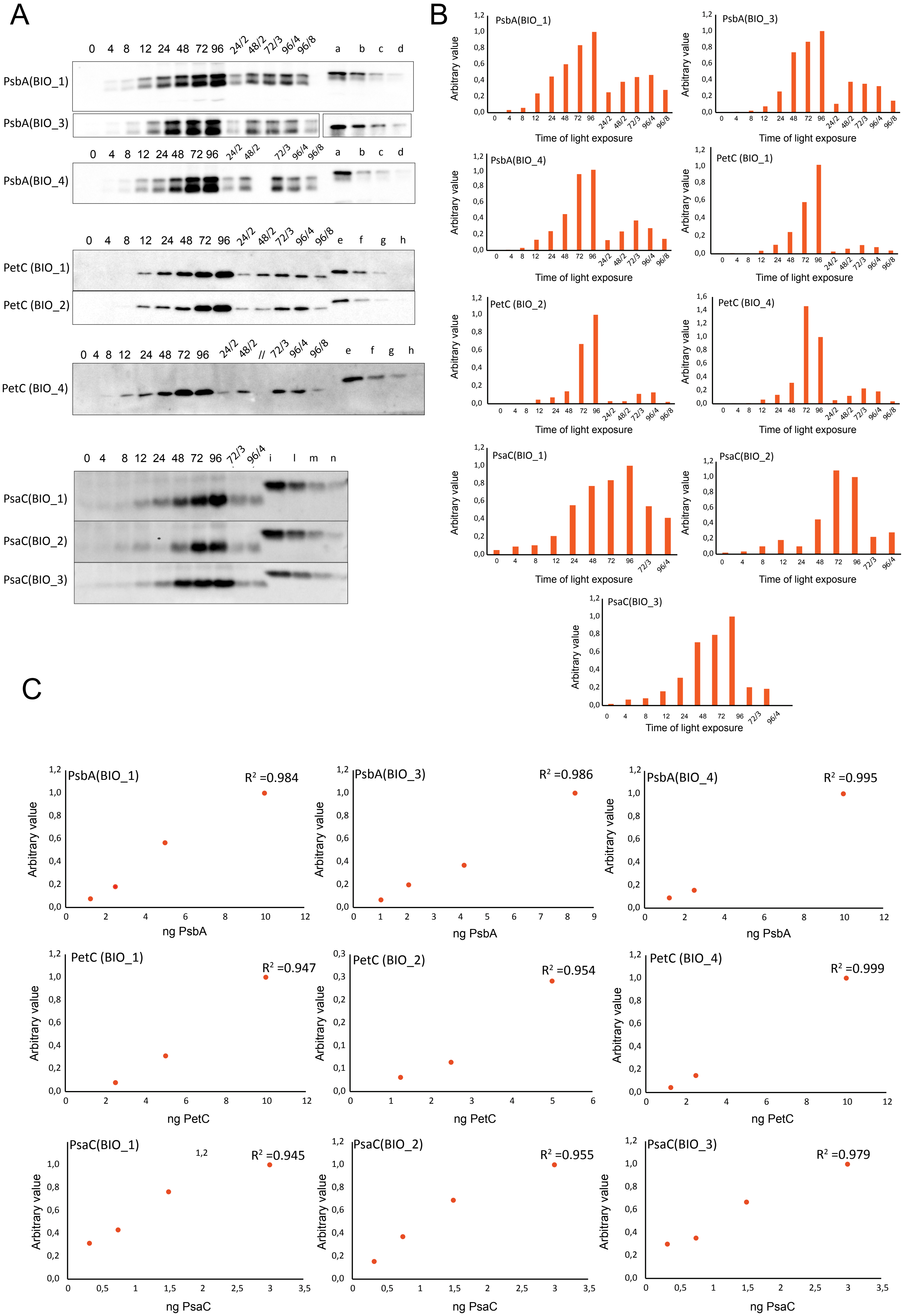

A
0 4 8 12 24 48 72 96 24/2 48/2 72/3 96/4 96/8 a b c d
PsbA(BIO_1)
PsbA(BIO_3)
PsbA(BIO_4)
0 4 8 12 24 48 72 96 24/2 48/2 72/3 96/4 96/8 e f g h
PetC (BIO_1)
PetC (BIO_2)
0 4 8 12 24 48 72 96 24/2 48/2 // 72/3 96/4 96/8 e f g h
PetC (BIO_4)
0 4 8 12 24 48 72 96 72/3 96/4 i l m n
PsaC(BIO_1)
PsaC(BIO_2)
PsaC(BIO_3)
B
PsbA(BIO_1)
PsbA(BIO_3)
PsbA(BIO_4)
PetC (BIO_1)
PetC (BIO_2)
PetC (BIO_4)
PsaC(BIO_1)
PsaC(BIO_2)
PsaC(BIO_3)
Arbitrary value
Time of light exposure
C
PsbA(BIO_1) $R^2$ =0.984
PsbA(BIO_3) $R^2$ =0.986
PsbA(BIO_4) $R^2$ =0.995
ng PsbA
PetC (BIO_1) $R^2$ =0.947
PetC (BIO_2) $R^2$ =0.954
PetC (BIO_4) $R^2$ =0.999
ng PetC
PsaC(BIO_1) $R^2$ =0.945
PsaC(BIO_2) $R^2$ =0.955
PsaC(BIO_3) $R^2$ =0.979
ng PsaC
Arbitrary value

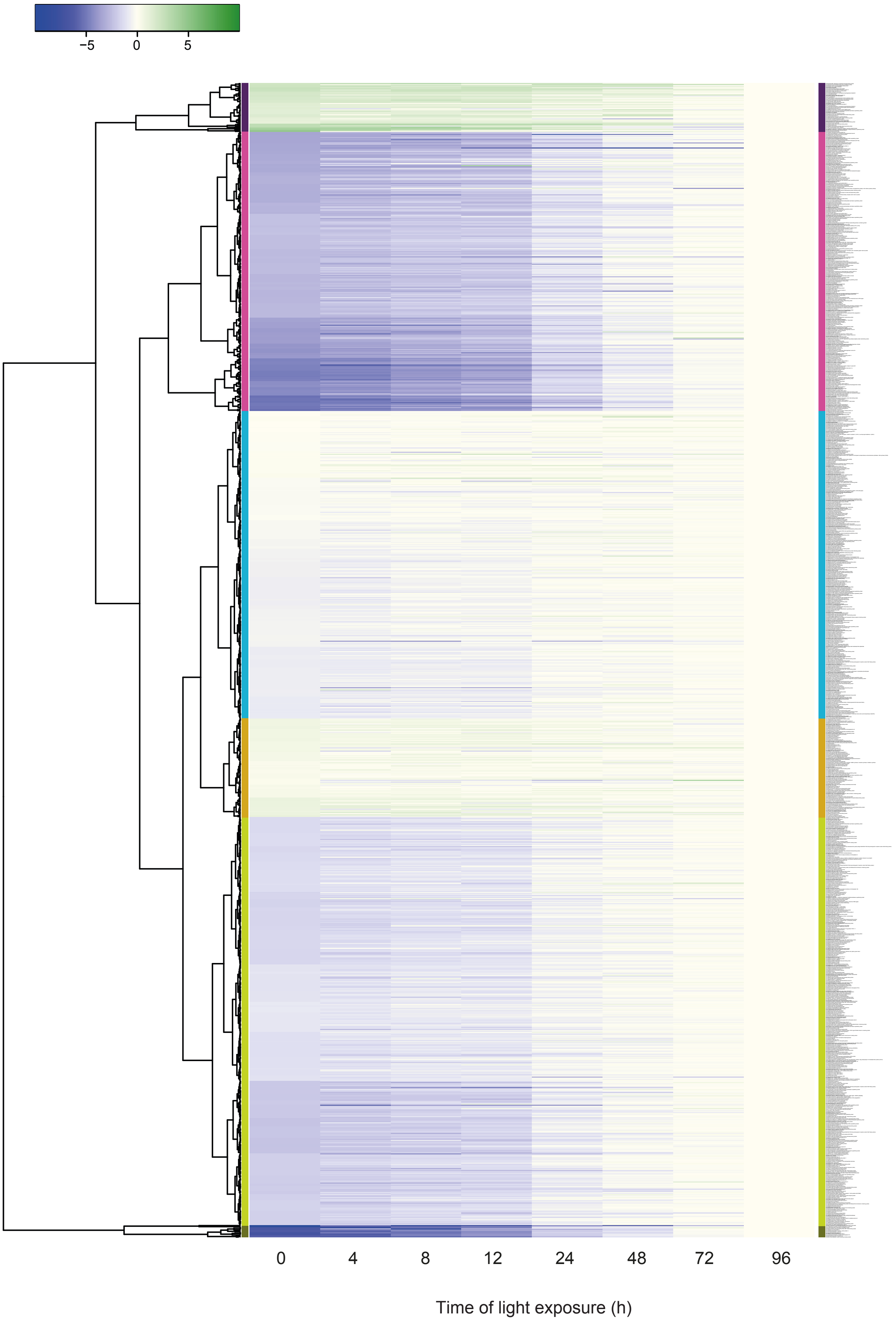
−5
0
5
0
4
8
12
24
48
72
96
Time of light exposure (h)

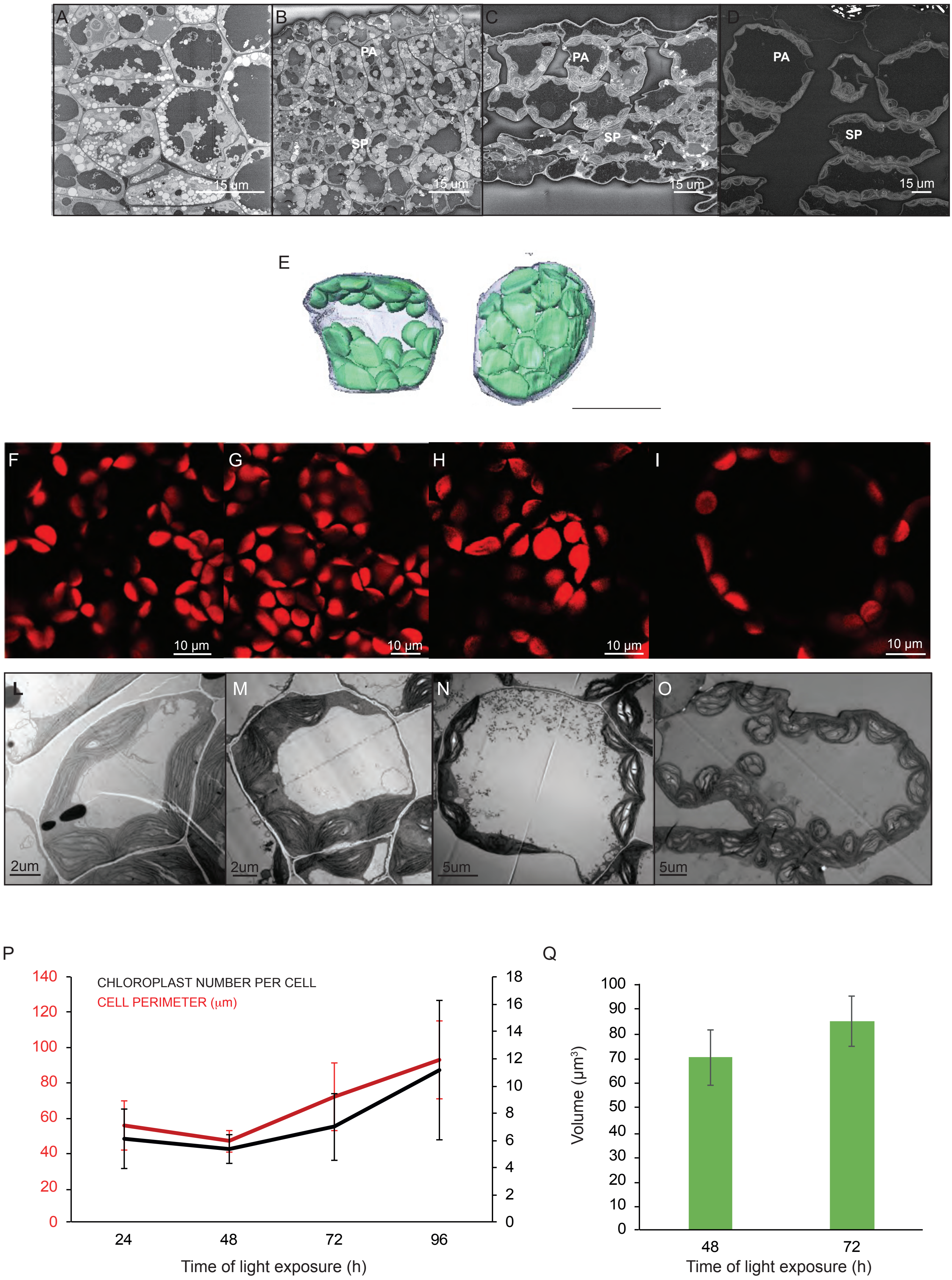

A
B
C
D
PA
SP
15 um
E
F
G
H
I
10 μm
L
M
N
O
2um
5um
P
CHLOROPLAST NUMBER PER CELL
CELL PERIMETER (μm)
Time of light exposure (h)
Q
Volume (μm³)
Time of light exposure (h)

A

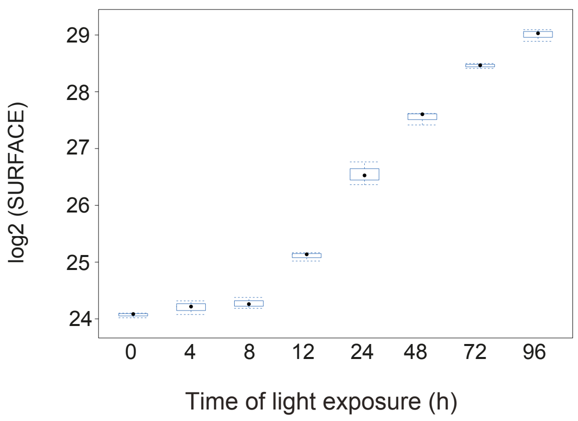


B

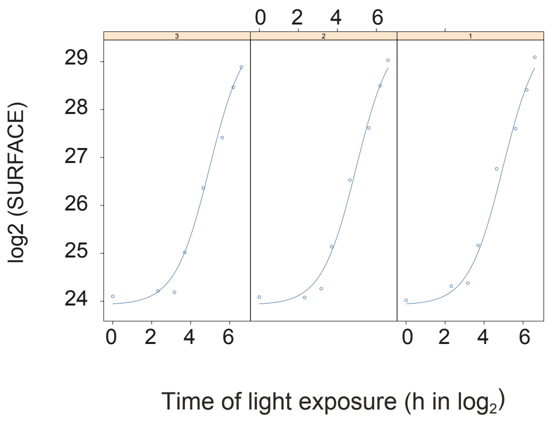


C

| Coefficient | Value | ± SE | t-Value | P value |
|---|---|---|---|---|
| *a* | 23.92 | ± 0.12 | 191.52 | 0 |
| *b* | 29.64 | ± 0.44 | 73.57 | 0 |
| *c* | 0.88 | ± 0.11 | 7.68 | 0 |
| *d* | 4.96 | ± 0.16 | 30.79 | 0 |

Equation 4 $$y = a + \frac{b - a}{1 + e^{(d - x)/c}}$$

A

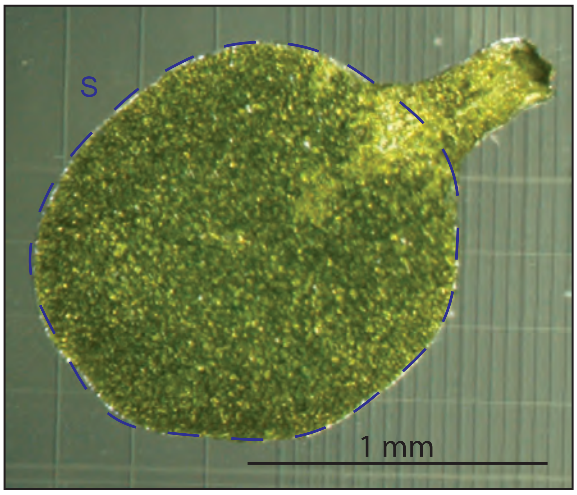


| TIME POINTS | T24 | T96 |
|---|---|---|
| Surface(S) | 0.33mm$^2$ SD +/-0.08 | 1.43mm$^2$ SD +/-0.07 |

B

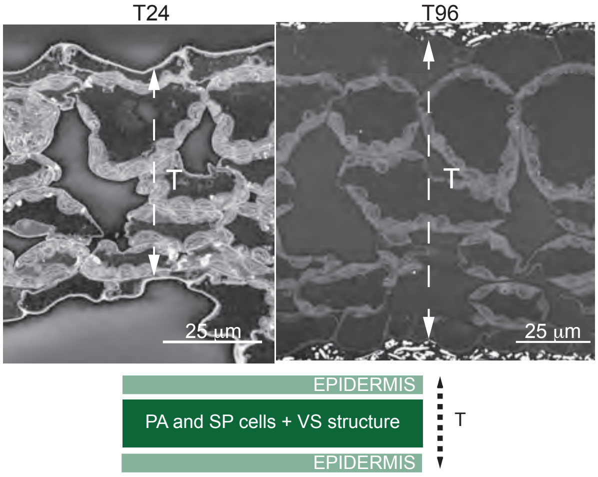


| TIME POINTS | T24 | T96 |
|---|---|---|
| Thickness (T) | 60 µm SD +/- 6 | 104 µm SD +/- 5 |

C

*Thickness (T)* * *Surface (S)* = COTYLEDON VOLUME

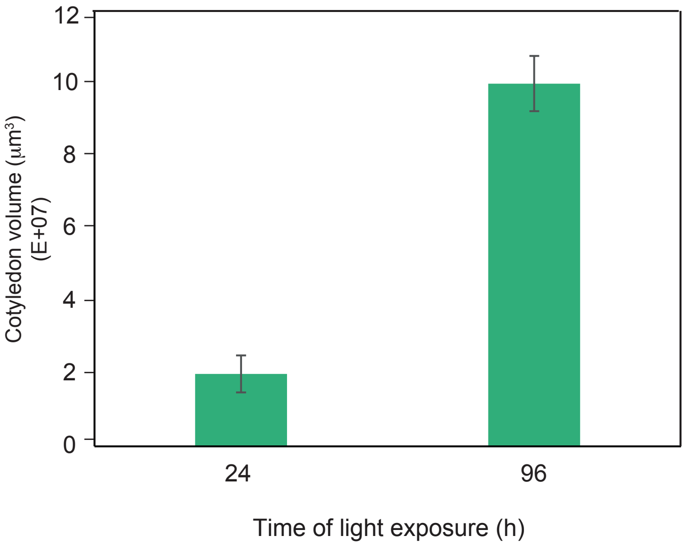


D

$$\frac{\text{COTYLEDON VOLUME} * 0.5}{\text{SINGLE CELL VOLUME}} \approx \text{NB.CELLS}$$

(Fig7B)

≈ 3000
